# Helical Intermediate Formation and Its Role in Amyloid of an Amphibian Antimicrobial Peptide^†^

**DOI:** 10.1101/2023.01.11.523545

**Authors:** Anup Kumar Prasad, Lisandra L. Martin, Ajay S. Panwar

## Abstract

Helical intermediates appear to be crucial in amyloid formation of several amyloidogenic peptides, including A*β*, that are implicated in different neurodegenerative diseases. Intermediate species have been reported to be more toxic than mature amyloid fibrils. Hence, the focus of the current work is to understand both structural and mechanistic role of intermediates in the early stages of amyloid self-assembly in amyloidogenic peptides. Molecular dynamics (MD) simulations and the adaptive biasing force (ABF) method were utilized to investigate structural changes that lead to amyloid formation in amphibian peptide uperin-3.5 (U3.5), an antimicrobial and amyloidogenic peptide. Microsecond time-scale MD simulations revealed that peptide aggregation, into *β*-sheet dominated aggregates, is centred on two important factors; evolution of *α*-helical intermediates and the critical role of local peptide concentration inside these aggregates. Electrostatic attraction between the oppositely charged aspartate (D) and arginine (R) residues located near the N-terminus induced hydrogen bonding resulting in formation of precursor 3_10_-helices close to the N-terminus. The 3_10_-helices transitioned into *α*-helices, thereby imparting partial helical conformations to the peptides. In the initial stages of aggregation, U3.5 peptides with amphipathic, partial helices aggregated to form small clusters of helical intermediates directed via hydrophobic interactions. These helices imparted stability to the helical intermediates, which promoted growth of clusters by further addition of peptides. This led to an increase in the local peptide concentration which enabled stronger peptide-peptide interactions and triggered a *β*-sheet transition in these aggregates. Thus, the study emphasized that stabilisation of peptide helical content may be crucial to the evolution of *β*-sheet-rich amyloid structures.

## 1 Introduction

More than fifty human diseases, called amyloidosis, are associated with deposition of amyloid proteins in tissues.^1,2^ Alzheimer’s disease, Parkinson’s disease, and type-2 diabetes are typical examples of amyloidosis in which *β*-sheet-rich amyloids can accumulate in affected organs.^3,4^ Amyloids are characterised by highly ordered *β*-sheet-rich aggregates which generally form by assembly into fibrillar structures of peptides and proteins. These peptide fibrils originate as intermediates that aggregate to form mature amyloid.^5^ However, these intermediate aggregates and protofibrils have been found to be more cytotoxic than mature amyloid fibrils and thus linked with pathological forms of amyloid diseases.^1,6^ The toxic nature of intermediate species has drawn the attention of researchers to investigate the mechanism of their formation and toxicity. Circular Dichroism (CD) and Infrared (IR) spectroscopic studies have revealed the formation of helical intermediates in the early stages of amyloid formation of A*β* peptide, *α*-synuclein, and islet amyloid polypeptide (IAPP, amylin).^7–11^ However, the mechanism of helical intermediate formation and its role in *β*-sheet-rich aggregation are still unknown.

Besides the pathogenesis of amyloids, they have been associated with vital biological functions, including antimicrobial activity, biofilm formation and cell adhesion.^4,12,13^ U3.5 is an amyloidogenic, antimicrobial peptide containing 17 amino acids (GVGDLIRKAVSVIKNIV-NH_2_), secreted on the skin of toadlets *Uperoleia mjobergii*.^14^ Interestingly, the U3.5 peptide has been called a chameleon peptide, showing cross-*α*/cross-*β* amyloid, under different conditions.^15^ Whether these chameleon-like properties of U3.5 are linked to its antimicrobial activity is still an open question. However, previous studies have shown that substitution of charged amino acids with hydrophobic residues or screening of charged residues by addition of salts increases the kinetics of U3.5 peptide aggregations.^16,17^ Bacterial membrane mimetic environments, including SDS (sodium dodecyl sulphate), DOPE (1,2-dioleoyl-sn-glycero-3-phosphoethanol-amine) and DOPG (1, 2-dioleoyl-sn-glycero-3-phospho-(1^′^-rac-glycerol)) and TFE (2,2,2-trifluoroethanol) solvents, have all been shown to induce the *α*-helical conformation in U3.5 peptides.^18,19^

Far UV CD spectra and ThT dye fluorescence assays have revealed the transient appearance of helical structures prior to the appearance of *β*-sheet-rich amyloids.^15,20^ The appearance of helices in these data, is associated with increased kinetics in peptide aggregation, indicating that there could be a role of helix formation in *β*-sheet-rich aggregation. Despite indications of helical intermediates in amyloid formation, it remains unclear how helical intermediates form and how they influence *β*-sheet-rich fibril formation. A pertinent question arises: whether the formation of helical intermediates is on-pathway of amyloid formation or if their formation is part of an off-pathway event. On-pathway helical intermediates can be good candidates in therapeutic development against amyloidosis. To understand the role of helical intermediates in *β*-sheet-rich aggregation of U3.5, it is important to investigate their content and stability at various stages of aggregation. For instance, helical intermediates were thought to be responsible for increased peptide aggregation in TFE, a known helical stabiliser. Whereas, peptide aggregation increased at low TFE concentrations (10 − 20%), it was inhibited at high TFE concentrations (≈ 50% v/v in water) where helices dominated.^20^

Furthermore, the role of amphipathic nature of U3.5 helices in promoting aggregation has rarely received comment. Intermediate species may arise during amyloid formation with heterogeneous and transient characteristics. The structures of short-lived (less than a few milliseconds) intermediate species and their role in peptide aggregation are challenging to investigate experimentally. Molecular dynamics (MD) simulations are ideally suited to investigate the emergence and interaction of diverse intermediates in amyloid formation at short time-scales. In addition, structural transitions within peptides occur at time-scales that are not easily accessible experimentally, but can be captured with MD simulations. Several MD simulation studies have been performed to investigate the oligomerization of peptides, elongation of fibrils and their stability.^21–23^ A*β* and *α*-synuclein have shown aggregation resistance by stabilization of helical conformers or lowering the population of partially structured helices.^24^ Hydrophobic interaction of termini (N-terminus of A*β*_42_ and C-terminus of *α*-synuclein) plays a crucial role in the stabilization of their central helical domains.^24,25^ MD simulation of A*β*_1−28_ showed the oligomerization, initiated by intermolecular *β*-sheets of N-terminal regions.^26^ In conjunction with free energy calculation methods, MD simulations have been used to examine energetics of structural transitions during aggregation of amyloidogenic peptides. In the current work, fully atomistic MD simulations were used to determine the role of helical intermediates in the aggregation of U3.5 peptides into *β*-sheet-rich structures.

## 2 Methods

### 2.1 Simulation Details

#### 2.1.1 Establishing a model for the U3.5 peptide

U3.5 peptide is known to adopt a random coil structure in pure water. The wild type peptide is C-terminal amidated at the final valine residue.^14^ A random coil structure of U3.5 with C-terminal amidation (generated using Visual Molecular Dynamics, VMD)^27^ was used as the starting structure in the MD simulations. In previous MD simulations of a single U3.5 peptide in solution, U3.5 conformations were found to contain largely random coil fractions.^17,19^ However, they also showed a significant amount of transient and partial helical content over hundreds of nanoseconds of simulation trajectories. Hence, a single peptide was simulated for 200 ns and several peptide conformations were randomly extracted from this trajectory. These were later used as representative peptide conformations for all simulations. Though the conformations remained predominantly random coil, several showed partial helical components in their secondary structure. Using these conformations, two simulation systems corresponding to different U3.5 concentrations were set up: (a) Unconstrained simulation: a simulation with twenty peptides (peptide concentration, *c*_*p*_ = 20 mM) and (b) Adaptive biasing force (ABF) simulation: a simulation to calculate the free energy landscape using four peptides (*c*_*p*_ = 30.75 mM).

The CHARMM36 force-field was used for fully-atomistic MD simulations in NAMD.^28^ Periodic boundary conditions were applied in all directions. A switching function for Lennard-Jones potential was used from 10 Å to 12 Å. Particle-mesh Ewald summation, with a grid spacing of 1 Å, was used to calculate electrostatic interactions.^29^ A time-step of 2 fs was used in the simulations. Temperature was maintained at 310 K using a Langevin thermostat with damping coefficient of 1 ps^−1^. A Nosé-Hoover Langevin piston, with a damping time constant of 100 fs, was used to maintain a 1 atm pressure in the system.^30,31^ The TIP3P^32^ water model was used for the solvent and an ionic strength of 0.15 M NaCl was maintained corresponding to physiological conditions. For the initial relaxation step, the systems were equilibrated for 300 ps in an NVT ensemble (*T* = 310 K), then for another 500 ps in an NPT ensemble with the peptide backbone restrained. The equilibrated systems were further simulated in an NVT ensemble.

#### 2.1.2 Unconstrained simulations

Twenty U3.5 peptides (*c*_*p*_ = 20 mM) were randomly distributed in a simulation box of volume 118 × 118 × 118 (Å^3^) using PACKMOL.^33^ After solvation with TIP3P water and addition of salt (NaCl) ions, the simulation box contained nearly 156,000 atoms. Three independent sets of the simulation were run for a duration of 2 *μ*s each.

#### 2.1.3 Adaptive biasing force simulation

The adaptive biasing force (ABF) method was used to estimate the potential energy surface corresponding to the aggregation pathway for a system of four peptides.^34^ Four U3.5 peptides were randomly distributed in a simulation box of 60 × 60 × 60 (Å^3^) with a minimum distance of 10 Å between any two peptides. The simulation box was solvated with TIP3P water at 0.15 mM NaCl salt concentration, resulting in a total of nearly 20,000 atoms. The ABF method was implemented in NAMD using the Colvars^35^ module. It was used to estimate the potential of mean force (PMF) as a function of two reaction coordinates. The two reaction coordinates were the radius of gyration (*R*_*g*_) and root-mean-square deviation (RMSD) of the peptides. A range of 0 – 20 Å for *R*_*g*_ and 0 – 30 Å for RMSD was chosen.

#### 2.1.4 Data analysis and plotting

VMD was used for visualization and some data analysis. The Matplotlib library of Python was used to plot the analysed data. The H-bond matrix was created using PConPy^36^ module of python. A sequence logo was generated using online server https://weblogo.berkeley.edu/.^37^ It should be noted that the term “helical” and “helices” have been used to refer collectively for *α*- and 3_10_-helices of peptides.

## 3 Results and discussion

### 3.1 Unconstrained simulations

Snapshots from 2 *μ*s long simulations of U3.5 peptides (*c*_*p*_ = 20 mM) are shown in Figure 1. As described in the Methods, several peptides were in partial helical conformations in the starting configuration (snapshot at (*t* = 0 ns). Initial and aggregated states for all three simulation sets are shown in Figure 1. In each set, U3.5 peptides were observed to simultaneously change secondary structure leading to the formation of various aggregates. The three trajectories showed significant helical (*α*-helix and 3_10_-helix) content during the aggregation process. The presence of helices can be seen in aggregates formed in the later stages of simulations in Figure S1. This observation suggested an important role for helical intermediates in the emergence of *β*-sheet-rich aggregates, at least, in the early stages of amyloidogenesis. Hence, a more detailed analysis of secondary structure evolution within these aggregates appeared to be a logical next step. At the end of 2 *μ*s simulations, at least two aggregates were observed in each trajectory, usually one much larger than the other. These were named using an alphanumeric code, where the letter S was used for the simulation set, A for aggregate number, and P for number of peptides in the aggregate. For example, the code S1A1P10 indicates (A)ggregate number 1 of (S)et 1 which contains 10 (P)eptides.

**Fig. 1.**
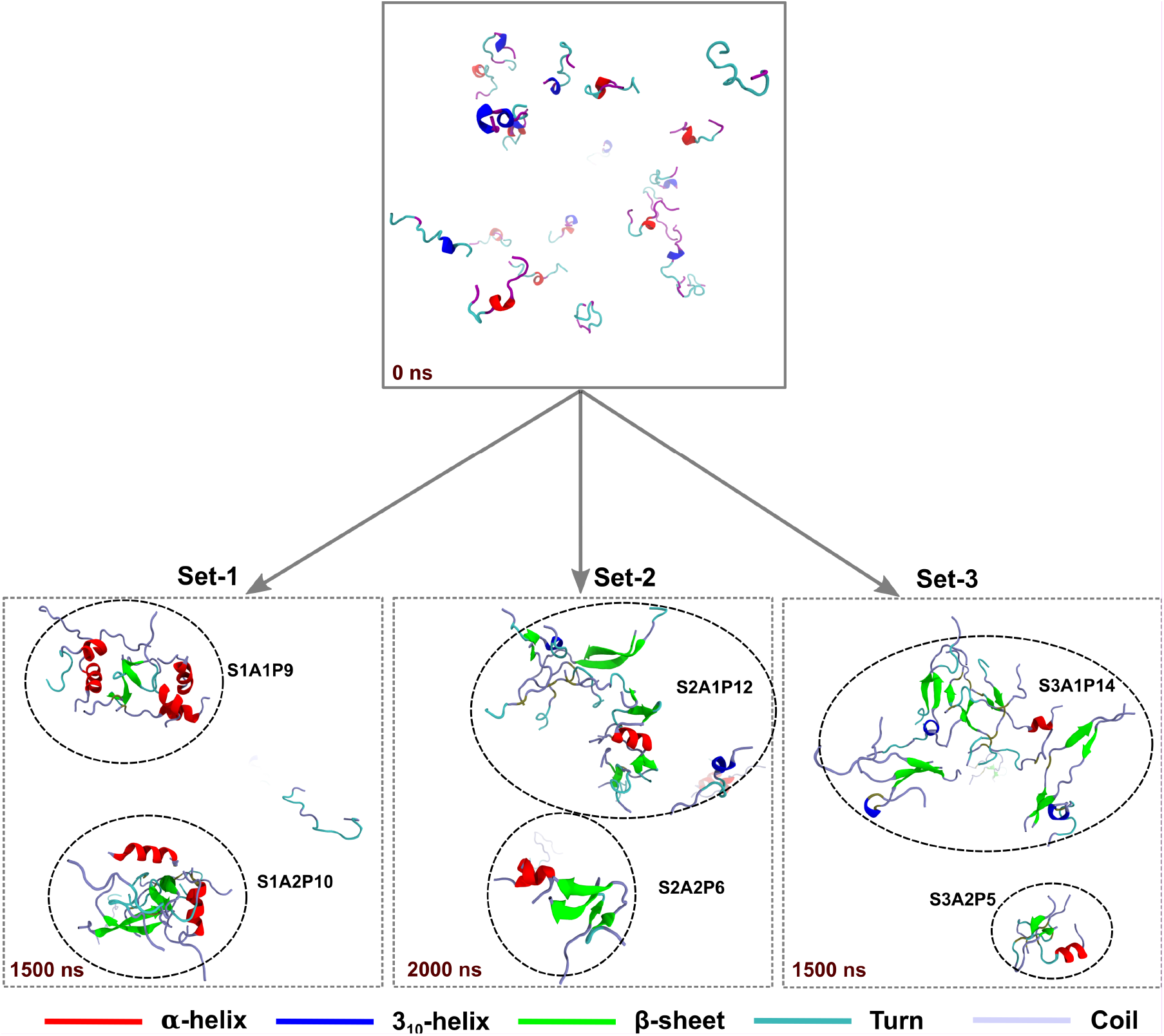
Snapshots of initial and later stages of the unconstrained simulations. The simulations, started with the peptides containing random coil and partial helical structures in unaggregated state, formed *β*-sheet-rich aggregates. Two aggregates were observed in each trajectory, highlighted by round circles.

The formation of aggregates appeared to follow diverse routes. In Set 1, a helical trimer was initially observed at ≈ 190 ns. Over the course of this simulation, the trimer interacted with six other peptides to form a stable aggregate of nine peptides (S1A2P9). The S3A1P14 was the largest aggregate formed among all three simulation sets that started to form at ≈ 225 ns in the simulation. Most aggregates were formed around 900 ns and remained stable throughout the simulations. However, peptide numbers within an aggregate remained dynamic for some aggregates. For instance, the evolution of S1A1P10 was characterized by dynamic attachment and detachment of peptides to the aggregate after ≈ 1100 ns of the simulation. This continued till the aggregate reached its final form at the end of 2 *μ*s. In addition, the S3A2P5 aggregate was completely disassembled at around 1.2 *μ*s. The dynamics of all these aggregates were analysed by quantification of the radius of gyration (*R*_*g*_) and root mean square distance (RMSD) for each aggregate (Figure S2). Every trajectory showed a continuous decrease in *R*_*g*_ within the first 900 ns corresponding to the formation of peptide clusters from initially isolated peptides. After aggregate formation, *R*_*g*_ usually fluctuated due to dynamic attachment and detachment of peptides to/from aggregates. Fluctuations in RMSD indicated structural changes in peptides within corresponding aggregates. The diversity among the aggregates, in the context of size and secondary structure composition, provides an indication of the polymorphism and heterogeneity that is often found in the early stages of amyloid formation.^38^ Previous CD and IR studies of amyloid formation have also revealed the appearance of helical structure in natively unfolded polypeptides, such as, in A*β* and *α*-synuclein.^39^ However, important questions concerning the significance of helical intermediates in toxicity or further development of amyloid fibrils remain unresolved. Another important question relates to the transition from helical intermediates to *β*-sheets during amyloidogenesis in these peptides.

#### 3.1.1 Role of helical peptides in *β*-sheet transitions

In our previous study, Ray *et al*. showed that U3.5 peptides with high helical stability had less propensity towards *β*-sheet aggregation.^17^ Interestingly, a small amount of *α*-helical stabilising co-solvent, TFE < 20% (v/v), was shown to accelerate U3.5 aggregation in water-TFE mixtures. However, higher concentrations of TFE > 20% (v/v) inhibited the peptide aggregation.^20^ Furthermore, MD simulations also showed that the presence of an SDS micelle, which stabilises the helical structure of U3.5 peptide, inhibits *β*-sheet aggregation by favouring intramolecular H-bonds over intermolecular H-bonds.^19^ These studies suggested that the amount of partial helical structure and its’ stability could play crucial roles in *β*-sheet transition, at least in the early stages of amyloid formation.

In order to understand the effect of peptide helical content and its stability in the formation of *β*-sheet-rich aggregates, a detailed analysis of the secondary structure evolution in aggregates was undertaken (Figure 2). This analysis showed significant variations in secondary structure evolution between the three simulation sets. The percentage of *β*-sheets gradually increased from almost nil to between 20% and 30% in all aggregates observed in all simulations. It is noteworthy that formation of *β*-sheets increased significantly after the formation of aggregates at 900 ns (Figure 2a). In contrast, helical content decreased in most aggregates with the exception of S1A2P9 and S2A1P12. These two aggregates showed an increase in helical contents till 1000 ns and 800 ns, respectively. However, in both aggregates, an increase in helical content in the initial stages was followed by helix to *β*-sheet transitions in the later stages of the simulations. Heat maps of the MD simulations (Figure S3) emphasised that secondary structure evolution during aggregation is characterised by the appearance of both transients as well as longer-lived stable helices (shown by short and long colour strips, respectively).

**Fig. 2.**
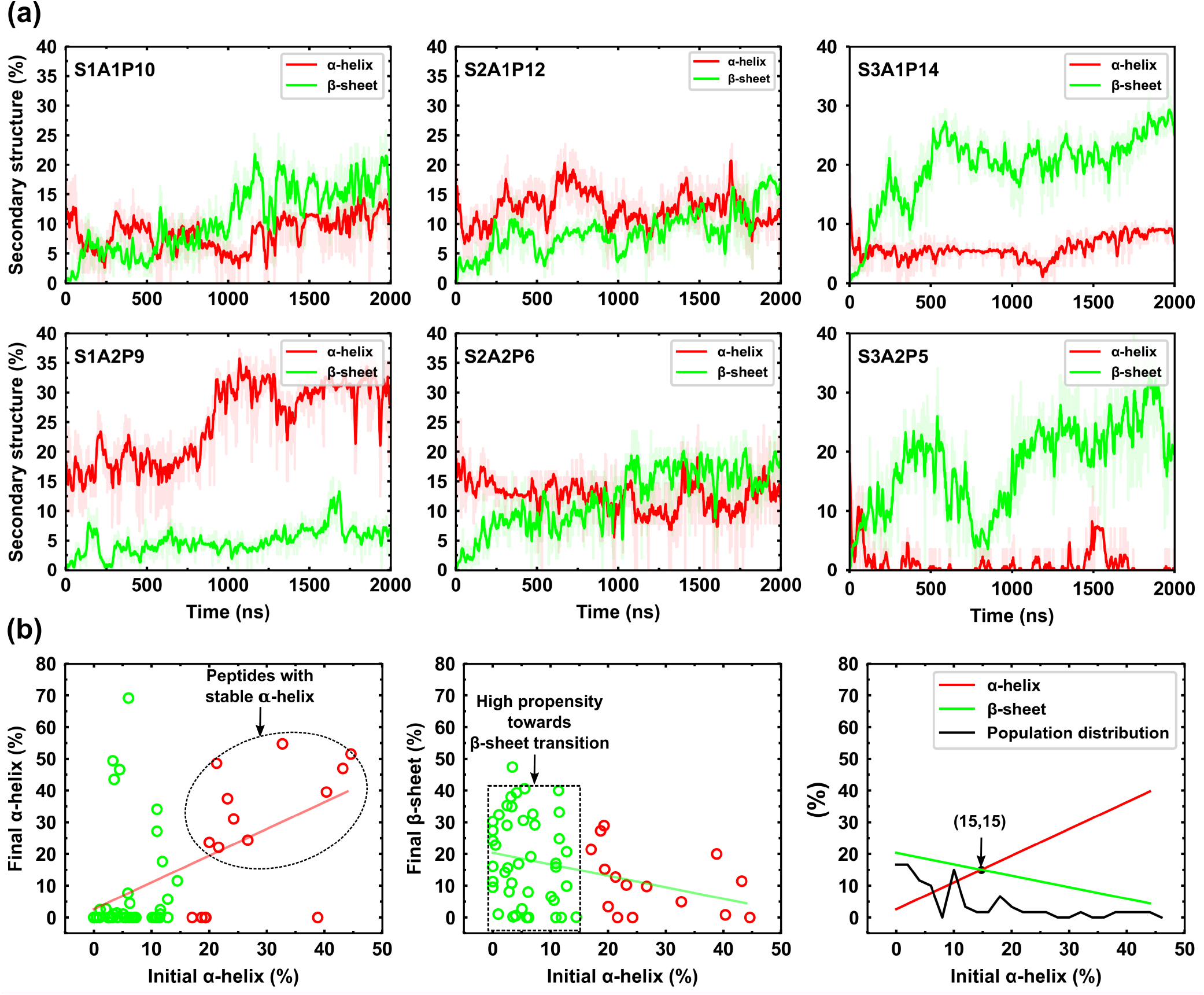
(a) Secondary structure evolution in the aggregates of unconstrained simulations. The percentage of *β*-sheet content gradually increased, significantly in the range where the helices decreased, with the exception of S1A2P9 and S2A1P12 aggregates. The S1A2P9 and S2A1P12 aggregates showed an increase in *α*-helices followed by *β*-sheet transition in peptides of aggregates. (b) Correlation of the peptide’s final secondary structures, at 2 μs, with the averaged helical content for the individual peptides of 0 to 50 ns of simulations. Each circle represents the peptide, initial helix < 15% (green circles) and initial helix > 15% (red circles). The last panel shows the intersection between the lines of best fit for data from the first and second panels. At the intersection point (15,15), the peptide has equal propensity toward *α*-helix and *β*-sheet transitions.

A correlation analysis between the final secondary structure composition (both *α*-helix and *β*-sheet) transitions with respect to the initial helical content of the peptides is shown in Figure 2(b). The stability of a secondary structure component was assumed if it remained stable for 50 ns. The initial helical content of a peptide was assumed to be its average helical content over the first 50 ns of the simulation. These results revealed that the initial helical content of peptides as well as their stability determined the fate of the final secondary structures of peptides within aggregates. Peptides with an initial helical content < 15% (shown by green circles) evolved mostly into *β*-sheets. In contrast, peptides with an initial *α*-helical content ≥ 15% (red circles) remained in *α*-helices till the end of the simulations. The cut-off 15% for initial helical content was based on the intersection value (15%) from lines of best fit of the structural correlation plots. It was found that peptides with low initial helical content showed a greater propensity towards *β*-sheet formation in aggregates. In contrast, peptides with higher initial helical content (≥ 15%) did not transition to *β*-sheet structures and remained stable in helical forms within the aggregates. A population distribution showed that 75% of the peptides were below the 15% cut-off value in which most of the peptides transitioned in *β*-sheets. The correlation analysis led to a more detailed examination of the molecular mechanism of *α*-helix to *β*-sheet transitions in peptides of aggregates.

#### 3.1.2 Influence of the local peptide concentration in aggregates

Secondary structure analysis of peptides in aggregates suggested that structural transitions were associated with an increase in local peptide concentration within these aggregates. For instance, respective helical contents in clusters S1A2P9 and S2A1P12 increased as the clusters became larger (i.e., the number of constituent peptides) and more compact (in size). This led to an increase in the local peptide concentration in proximity to the aggregates. Interestingly, the increase in local peptide concentration was followed by helix to *β*-sheet transitions within the compact aggregates. Like most AMPs, the U3.5 peptide forms an amphipathic *α*-helix that played a crucial role in peptide aggregation. These events were most dramatically observed in the aggregate S1A2P9. In the initial stages of S1A2P9 aggregate formation (see Figure 3), three peptides in partial helical conformations were found to assemble into a stable helical trimer at around 190 ns (conformer B). All three peptides adopted amphipathic helical conformations and associated in a way that the hydrophobic surfaces of the individual helices (shown in yellow) were buried inside the aggregate. This left the hydrophilic surface (red) exposed to the solvent facing outwards. Interestingly, the helical trimer remained stable throughout the simulation and seemed to serve as a “nucleus” for subsequent aggregation events. In our previous study, Ray *et al*. showed that U3.5 peptides in amphipathic partial helical conformations aggregated to form small clusters, burying their hydrophobic residues in the aggregated core. They argued that peptides with partial helical conformations might provide an easier route to stable aggregates, in contrast to those with random coil peptides which are prone to thermal fluctuations and disruption.^17^

**Fig. 3.**
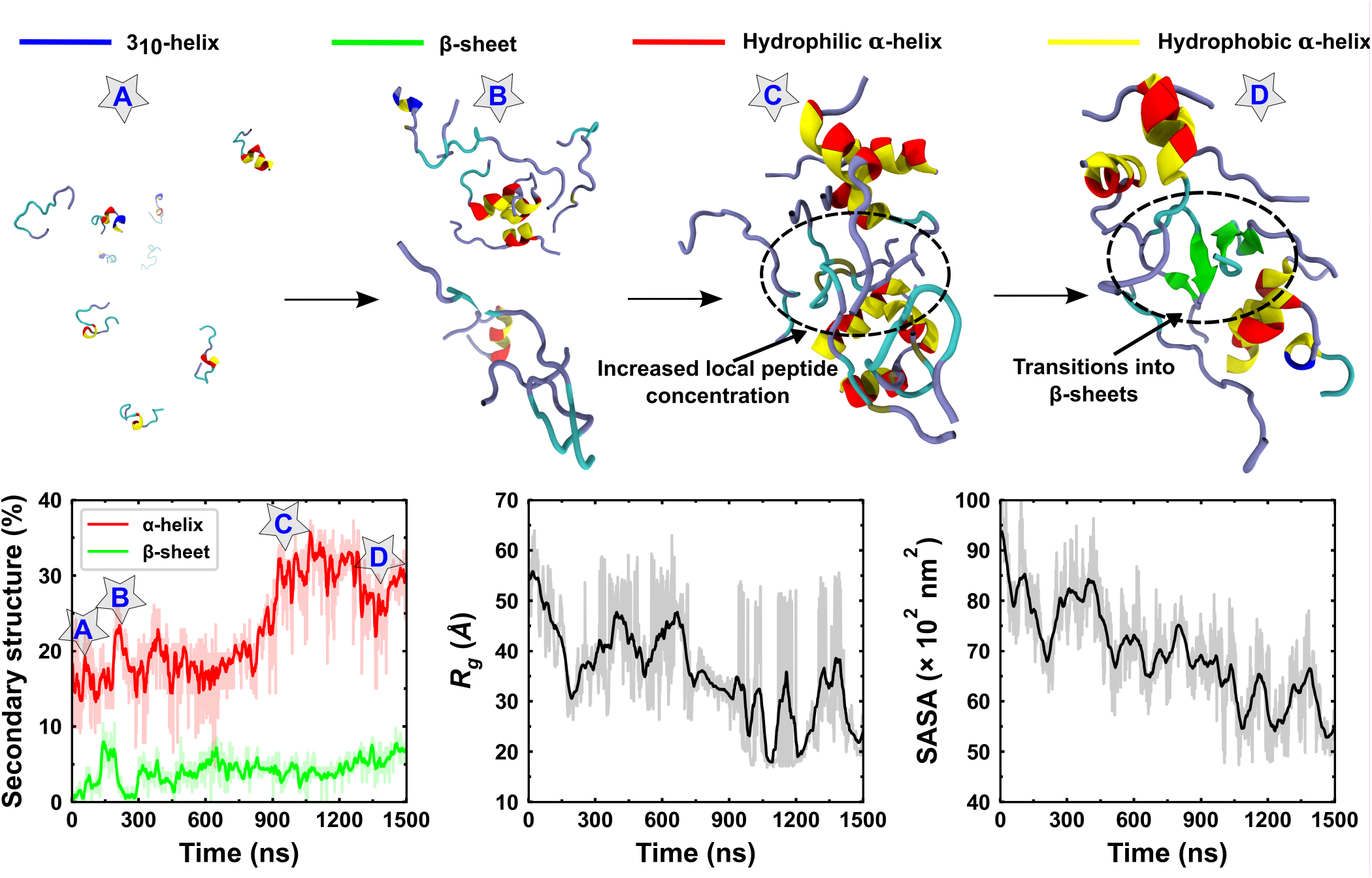
The snapshots of the S1A2P9 aggregate corresponding to different stages of helical intermediate formation. The peptides containing the random coil and partial helical structure were initially in an unaggregated state (conformer A) that formed a stable helical trimer associated with other peptides at 190 ns (conformer B). Conformer C shows the increased local peptide concentration dominated by the helical intermediate which facilitates the *β*-sheet transitions in conformer D. The plots of secondary structure, *R*_*g*_ and, RMSD show the aggregate evolution. The *α*-helix content of the aggregate was significantly increased from 600 to 1000 ns which leads to the formation of the helical intermediate with increased local peptide concentration (Conformer C). In transition from conformer C to D, the *α*-helical content decreased with incremental increase of *β*-sheet. The radius of gyration (*R*_*g*_) and solvent accessible surface area (SASA) gradually decreased over time, that indicated the progression of aggregate compactness.

In the case of S1A2P9, the helical trimer attracted other individual peptides to adsorb onto the trimer, thus making the aggregate even larger. This process led to an increase in the local peptide concentration in the region around the aggregate and triggered a further increase in *α*-helix components of the aggregate. Between 750 ns and 1100 ns, the helical content in S1A2P9 increased from ≈ 20% to ≈ 35%. During this process, both the radius of gyration (*R*_*g*_) of aggregate and the solvent accessible surface area (SASA) of hydrophobic residues reduced significantly, indicating formation of a compact aggregate with hydrophobic residues buried deep inside the aggregate. This can be understood as a direct consequence of large-scale secondary structure transitions into *α*-helices that enabled peptides to pack more compactly inside the aggregate core. At 1100 ns, the aggregate was in a “sandwich-like” structure, where random coil segments of peptides were sandwiched between two large helix-dominated domains. The confinement of random coil segments between the surrounding helical domains resulted in an increased local peptide concentration with enhanced peptide-peptide interactions. In addition, the helical domains provided additional stability to the random coil segments, which assisted the transition of random coils to *β*-sheets in that region.

Though, aggregate S2A1P12 did not display dramatic increases in helical content during its formation, secondary structure transitions within S2A1P12 appeared to be qualitatively similar to those observed for S1A2P9. S2A1P12 also formed a *β*-sheet-rich aggregate (Figure S5), resulting from the association of amphipathic helices and increase in local peptide concentration. Until 700 ns, the helical content in S2A1P12 increased two-fold (conformer B). S2A1P12 formed a sandwich-like structure which facilitated peptide-peptide interactions due to an increase in local peptide concentration (conformations C and D). During the transition to *β*-sheets, both aggregates showed significant losses in their helical contents. Visual inspection revealed some peptides in these aggregates expanded *β*-sheet segments toward the helical segments leading to uncoiling of the helices. These results provide novel insight into the role of an unstable partial helix in increasing local peptide concentration and subsequent *β*-sheet formation in aggregates. Thus, a likely route for the formation of *β*-sheet-rich aggregates appeared to emerge from the simulations, involving the following steps: (i) initial aggregation of amphipathic helices, (ii) further incorporation of peptides through hydrophobic interactions, (iii) increase in helical content, (iv) compaction and increase in local peptide concentration and (v) transition to *β*-sheets.

### 3.2 ABF simulation

The formation of fibrillar amyloids from natively unfolded peptides, including U3.5, is complex and may involve a number of intermediate states.^40,41^ The MD simulations discussed above highlighted the role of helical intermediates in the early stages of aggregation. However, a simulation with twenty peptides corresponds to a large number of degrees of freedom which require extended trajectories to unravel all possible transitions. For example, certain conformations may remain trapped in kinetically limited states over computationally-limited timescales. Therefore, the ABF method (an enhanced sampling technique) was utilized, for a smaller system of four peptides, to elucidate aggregation pathways for the U3.5 peptide.

An ABF simulation with four U3.5 peptides was carried out for a duration of 3 *μ*s. At the start of the simulation, the peptides existed in random coil conformations and were separated from each other by a distance of 10 Å. The potential mean of force (PMF) for the aggregation process was computed as a function of two reaction coordinates; the radius of gyration (*R*_*g*_) of the four peptides and their root-mean-square deviation (RMSD) (calculated with respect to the starting state). Consistent with the unconstrained simulations, the ABF simulation also showed structural transitions from random coils to intermediate *α*-helices which, in turn, evolved into *β*-sheets. The PMF from the ABF simulation is shown in (Figure 4). Representative conformer snapshots corresponding to important transitions during the aggregation process are also shown. The PMF plot shows a well-defined free energy landscape with a global minimum corresponding to a *β*-sheet-rich aggregate. Conformational analysis of the simulation unravelled the aggregation pathway of U3.5 peptides. Six representative conformers were identified from the simulation and were named T1 to T6. The suffix numbers indicate the chronology of aggregation events and also decreasing energy levels from T1 to T6. Structural snapshots of these conformers were captured and their secondary structure components were analysed. Aggregation of the four peptides started with the highest energy conformer T1, which consisted of isolated, random coil peptides. During the aggregation process, T1 underwent transitions through different intermediate states to form a compact *β*-sheet-rich aggregate T6 (lowest energy conformer).

**Fig. 4.**
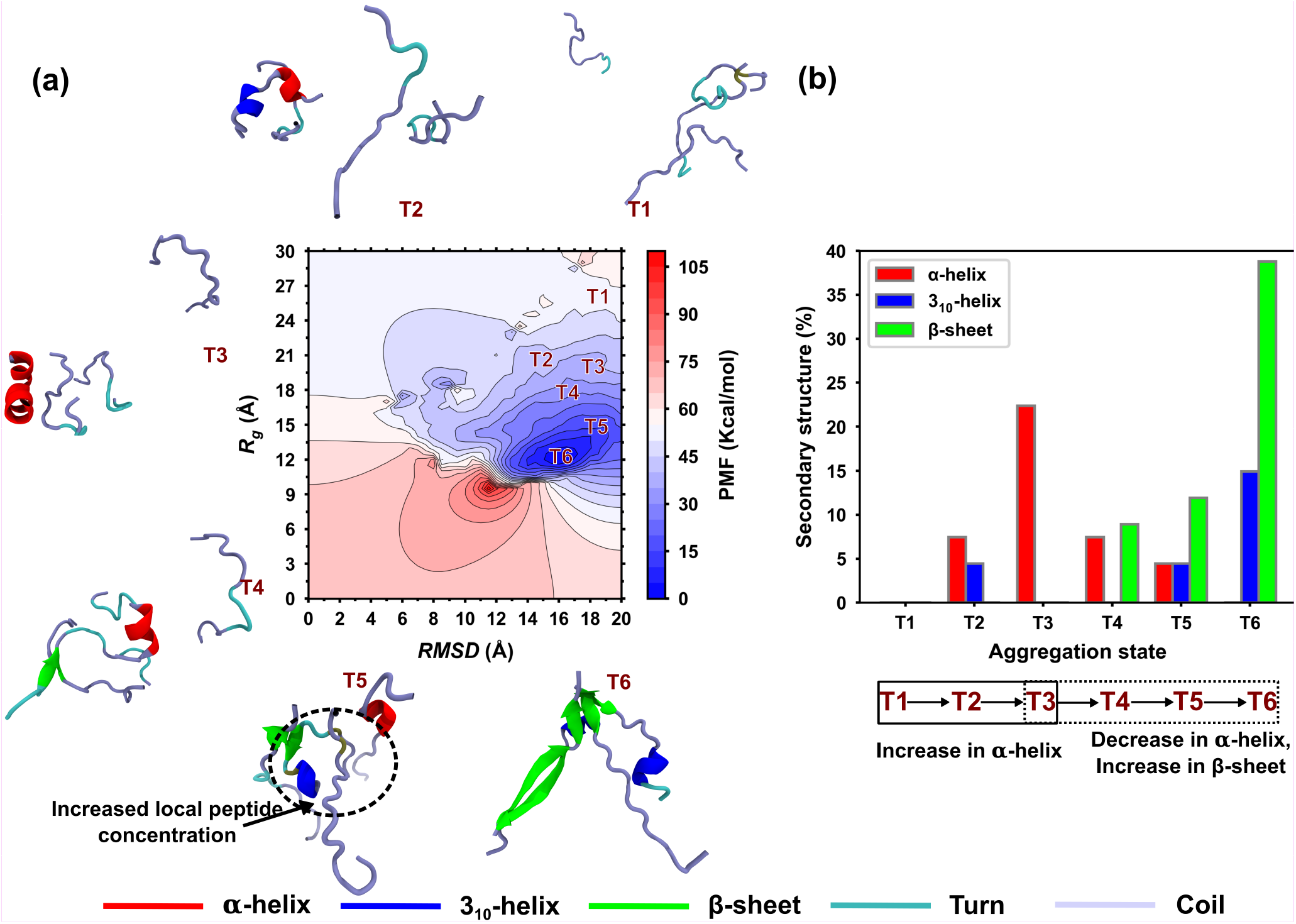
(a) The free energy landscape is shown as a function of *R*_*g*_ and RMSD reaction coordinates obtained from the ABF simulation. The PMF profiles of conformers, corresponding to different stages of aggregation, show the progression of aggregate formations from top to bottom of the energy well. (b) The secondary structure analysis of conformers showed the continuous increment of helices in helical intermediate formation (T1→T3) that further transformed into *β*-sheets (T4→T6).

In the aggregation sequence T1→T2→T3, one of the four peptides transformed into a partial *α*-helix and started associating with the other three peptides. The association of the helical peptide with other random coiled peptides triggered formation of the cluster T2. Subsequently, the *α*-helical content of the peptide cluster increased three-fold in the T2→T3 transition to form a more compact aggregate structure. A further increase in local peptide concentration was observed in T4, which facilitated increased peptide-peptide interaction, and resulted in the initiation of *β*-sheets formation in the aggregate. Finally, the peptide aggregation sequence T4→T5→T6 was characterised by a gradual decline in the *α*-helical content, which completely disappeared in T6. This contrasted with a gradual increase in *β*-sheet content, which reached a maximum in T6. Figure 4b shows the increase in *α*-helix for aggregation sequence T1→T3, and the subsequent decrease in *α*-helix for the aggregation sequence T4→T6. The *α*-helix to *β*-sheet transition for T4→T6 is also clearly evident from Figure 4b. More importantly, the PMF clearly showed that the energy decreased for the aggregation sequence T1 to T6. Whereas, T1 was at top of the energy well, T6 existed at the bottom of energy well and represented the most stable state in the simulation. This supported the hypothesis that helical intermediates lie along the pathway to formation of *β*-sheet-rich aggregates for U3.5 peptide. The transition from random coil, isolated peptides (T1) to a *β*-sheet-rich aggregate (T6) occurred through an *α*-helix rich intermediate state (T3).

These results show the significance of the helical intermediates for peptide-peptide interactions in order to form *β*-sheet-rich aggregates from unstructured peptides. However, an important question still remained unanswered: how is *α*-helix formation initiated in peptide clusters and why does it disappear during the transition to *β*-sheet-rich aggregates? To address these questions, we further analysed the 3_10_-helices, residual interactions and H-bonding in peptides.

#### 3.2.1 3_10_-helix as a conformational transition state

The 3_10_-helix is the fourth most common motif found in peptides and proteins, after the *α*-helix, *β*-sheet and turns. It is reported to be a short-length motif, yet it plays a significant role in structural transitions in peptides.^42,43^ Interestingly, several short (about 4 residues) 3_10_ helix structures were observed in all U3.5 trajectories (including unconstrained and ABF simulations) investigated here. These 3_10_-helices were found to be temporally unstable and were mostly associated with the formation of primordial *α*-helices. They vanished later in simulation trajectories as *α*-helices became stable and grew in content. The emergence and decay of the 3_10_-helices in the ABF simulation are analysed in greater detail in the heat map in Figure 5(a). Though, 3_10_-helices emerged in all four peptides over the entire trajectory, they were primarily observed in peptides P1 and P2. Their occurrence was strongly correlated with the occurrence of *α*-helical structures in the peptides. As a result, 3_10_-helices were observed in P1 and P2, where a majority of *α*-helices formed during aggregation (T1→T3) and later disappeared as they transformed to *β*-sheets (T4→T6). Figure 4(b) and 5(a) suggest that 3_10_-helices acted as transition states for both coil to *α*-helix and *α*-helix to *β*-sheet transitions. In Figure 5(a), P1 acquired an *α*-helical structure via 3_10_-helix starting at around 500 ns. At a later stage, a 3_10_-helix transition state is again observed in P1 during a *α*-helix to *β*-sheet transition (denoted by a dashed circle and arrow around 1400 ns). Similarly, 3_10_-helices were also observed in all sets of the unconstrained twenty peptide simulations throughout the trajectories (Figure S3). The occurrence of 3_10_-helices close to critical structural transitions indicates that they play a significant transitional role, in both coil to *α*-helix and *α*-helix to *β*-sheet transitions. Thus, 3_10_-helices are crucial not only to the evolution and stability of intermediate *α*-helical states in U3.5 aggregates, but also to the evolution of *β*-sheets from intermediate *α*-helical states. However, it should be noted that a few random coiled peptides directly transformed into *β*-sheet structures without undergoing transitions via 3_10_-helices.

**Fig. 5.**
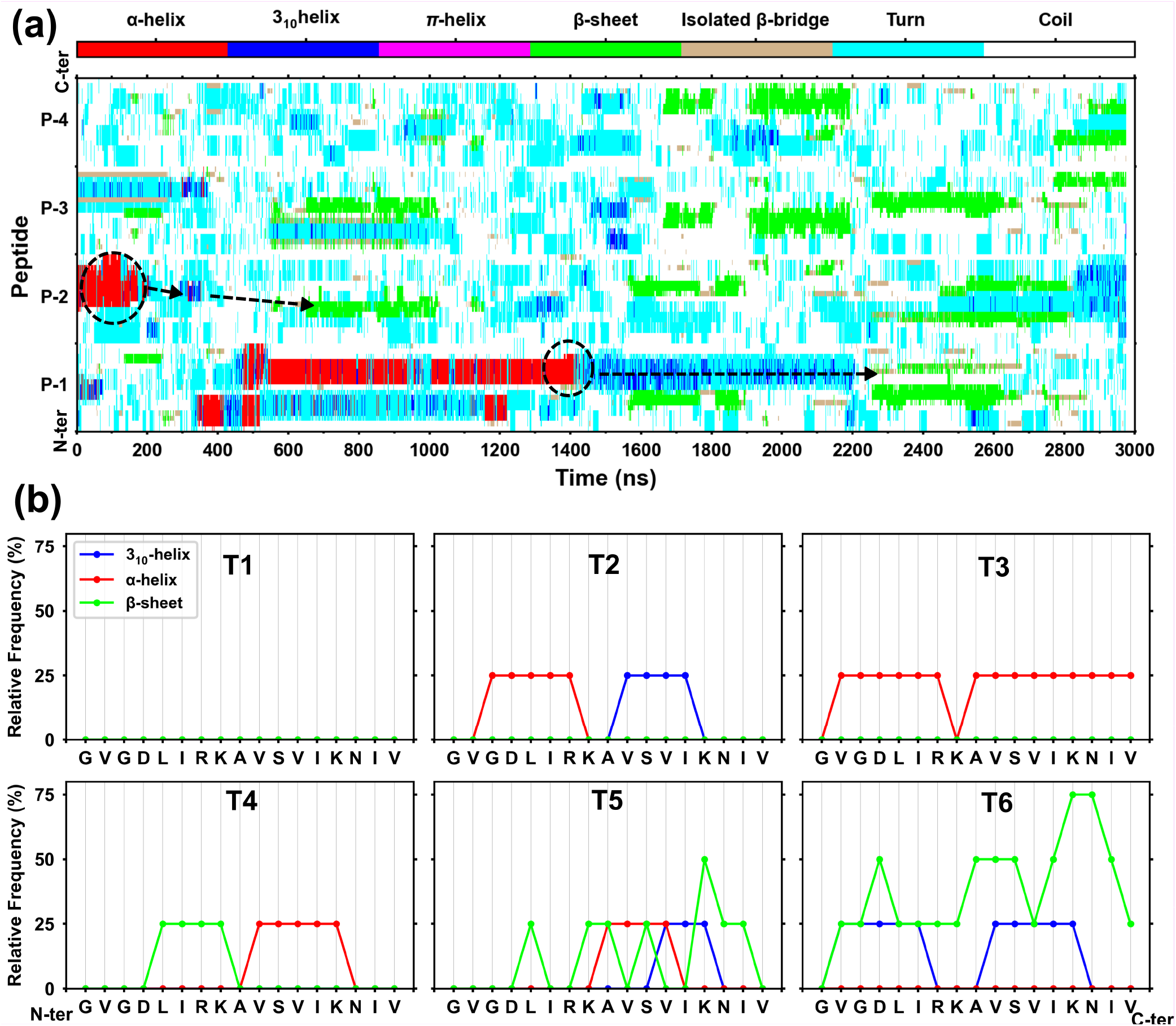
(a) Heat map of secondary structure evolution obtained from the ABF simulation. The transitions of *α*-helix to *β*-sheet occurred in the peptides P1 and P2 via 3_10_-helices. (b) Relative frequencies of secondary structure components are shown along peptide length for all representative conformers of aggregation. The *α*-helix formation started near N-terminus that further extended toward C-terminus by mean of 3_10_-helices (conformers T2→T3). The transitions of *α*-helix to *β*-sheet occurred via 3_10_-helices (conformers T4→T6).

#### 3.2.2 Residual interactions in helix to *β*-sheet transition

To investigate the role of residues in *β*-sheet-rich aggregation, residue-wise secondary structure and H-bond distributions along the residues were analysed. Figure 5(b) shows the distribution of relative frequency of secondary structure occurrence across the residues for all six conformers from the ABF simulation in Figure 4. In T2, an *α*-helix formed in the 3-DLIR-7 segment of U3.5, which later extended towards the C-terminus in T3 via 3_10_-helix transition (first observed in T2). An *α*-helix to *β*-sheet transition was observed from T3 to T4 as a *β*-sheet appeared in T4 toward the N-terminus of the peptide. From T4 to T5, *β*-sheets further extended towards the C-termini of the peptides. Concurrently, the *α*-helix content was much reduced in T5 and 3_10_-helix components were observed indicating their transitionary role in *α*-helix to *β*-sheet transition. The maximum *β*-sheet content was observed in T6 with the highest relative frequency of occurrence (nearly 75%) toward the C-terminus of the peptides. This suggested a strong preference for the C-terminus in the initiation of *β*-sheets in U3.5 aggregates.

The time evolution of various secondary structure components along the length (residue-wise distribution) was also investigated for the 2 *μ*s long unconstrained simulations. Figure 6 shows the residue-wise frequency distribution of various secondary structure components, averaged over all three trajectories of the unconstrained simulations and also over intervals of 400 ns. Interestingly, 3_10_-helices were induced in 4-DLIR-7 and 12-VIKN-15 segments of U3.5 peptide throughout the simulations (Figure 6a). The relative frequency of 3_10_-helix occurrence at 4-DLIR-7 was approximately double compared to its occurrence in the 12-VIKN-15 segment, although both were low. The N-terminus showed a greater propensity towards *α*-helix formation, in sharp contrast with the C-terminus where *α*-helices were virtually absent (Figure 6b). This observation further validated the strong link between the occurrence of 3_10_- and *α*-helices, since 3_10_-helices were shown to be in greater proportion toward the N-terminus (Figure 6a). However, the 3_10_-helices were found at both 4-DLIR-7 and 12-VIKN-15 segments, whereas the *α*-helix registered a maximum frequency near the central segments. In contrast, with the occurrence of *α*-helices near the N-terminus, the *β*-sheet formation occurred more across C-terminus (Figure 6d), although the frequency was similar to that observed for the *α*-helices. Interestingly, a clear and sharp maximum in *β*-bridges towards the C-terminus could be a precursor to the *β*-sheet formation. In addition, and conversely to the *α*-helix pattern (Figure 6b), *β*-sheets were absent at the N-terminus of the U3.5 peptides (Figure 6d).

**Fig. 6.**
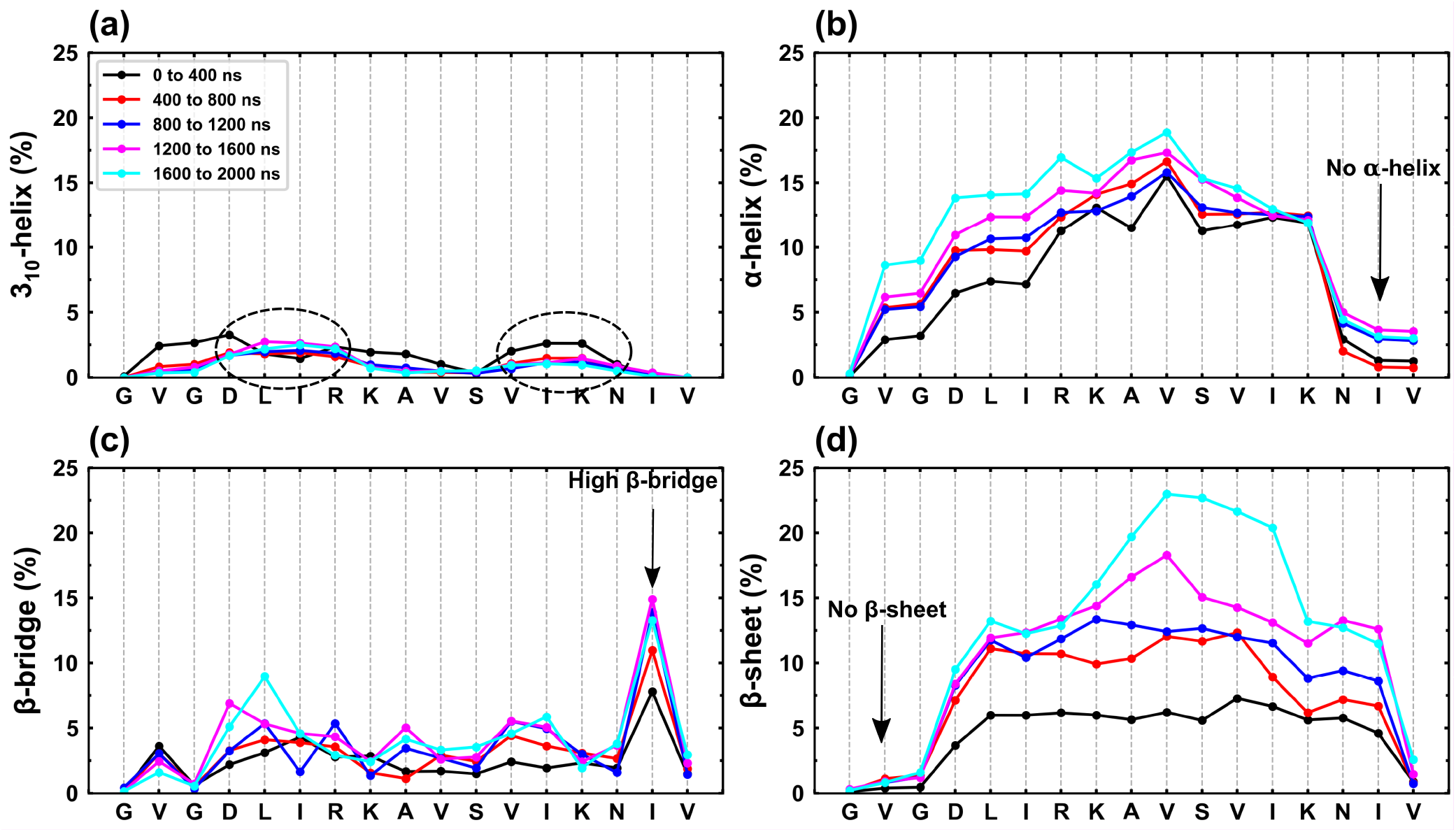
Relative frequencies (averaged) of secondary structure components are shown along the peptide length. Secondary structure occurrence was calculated for all trajectories of unconstrained simulations over the mentioned period of time. The 3_10_-helices were formed predominantly in 4-DLIR-7 and 12-VIKN-15 segments of U3.5 peptides.

This residue-wise analysis of secondary structure distribution along the U3.5 peptide provided crucial insight into *α*-helices being induced predominantly toward the N-terminus of the peptide. Hydrophobic interactions between amphipathic helices could also lead to aggregation of peptides resulting in the formation of helical intermediates in the course of aggregation. Once the aggregates were formed, peptide-peptide interactions towards C-terminus led to *β*-sheets transitions in peptides. Furthermore, the inter-peptide contacts map of conformers (Figure S6) showed strong peptide-peptide interactions towards N-terminus during the formation of helical intermediates (T1→T2). However, during the transition of the helical intermediate to a *β*-sheet aggregate (T4→T6), peptide-peptide interactions shifted toward the C-terminus of the peptides.

#### 3.2.3 Hydrogen bond analysis

The secondary structure of peptides and proteins is essentially formed by regular inter-molecular and intra-molecular hydrogen bonding of backbone amide groups. Hydrogen bond (H-bond) networks are also used to explore the secondary structure evolution in proteins.^44^ In a similar way, the evolution of H-bond networks from one conformer to another was analysed to understand structural transition both leading to and emanating from, the helical intermediate in the ABF simulation. Figure 7 shows H-bond networks between the four peptides in the ABF simulation, P1 to P4, using a grid-based “contact map”, where the black grid represents each peptide, and the finer divisions of the amino acid residues. Intra-peptide and inter-peptide H-bonds are represented by red and blue grids, respectively. The change in H-bonding pattern is clearly evident as the system transitions through different secondary structure components at various stages of the aggregation process. The 3_10_-helix and *α*-helix show 2 and 3 empty grids in these plots, respectively, because H-bonds form between *i*^*th*^ and (*i* + 3)^*th*^ (3_10_-helix) or *i*^*th*^ and (*i* + 4)^*th*^ (*α*-helix) amino acid residues of each peptide. For the sequence, T1→T3, P1 transitioned from random coil to helix to form a helical intermediate. Intra-molecular H-bonding (intra-HB) interactions at the N-terminal of P1 in T1 led to the formation of a 3_10_-helix that further transformed into *α*-helix in T3.

**Fig. 7.**
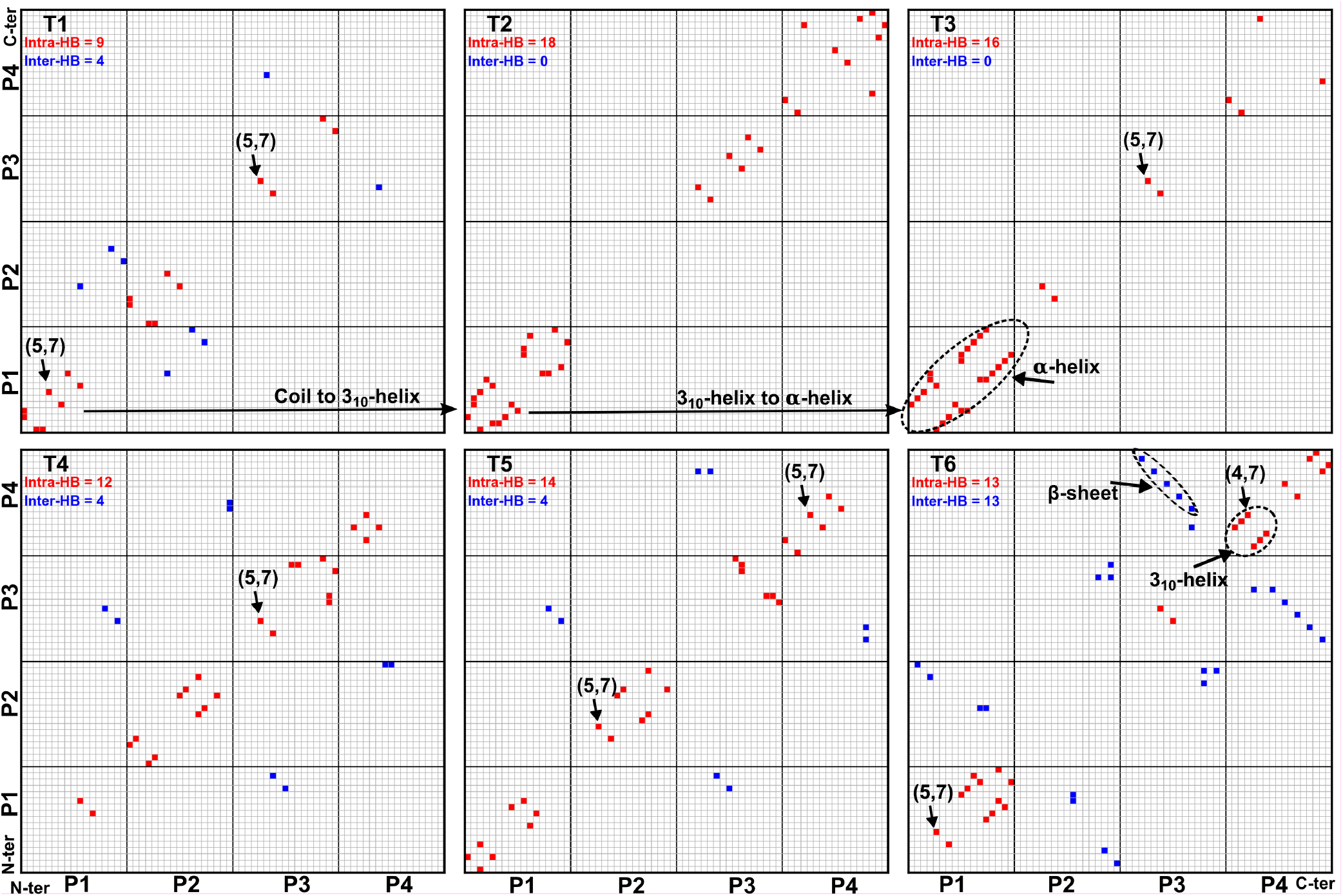
Hydrogen bond networks of the conformers are depicted, where each small grid represents a single residue. Intrapeptide and interpeptide hydrogen bonds are shown in red and blue grids respectively corresponding to residues of the peptides. The evolution of the hydrogen bond patterns in structural transitions can be seen in T1→T3 and T4→T6 conformers. The 3_10_/*α*-helix shows 2/3 empty grids in network, because H-bonds form between *i*^*th*^ and (*i* + 3)^*th*^/(*i* + 4)^*th*^ residues of peptide. The antiparallel *β*-sheet formed between peptide P3 and P4 in conformer T6. Most of the peptides formed H-bonds between L(5) and R(7) amino acids that are shown at (5,7) coordinate in the Figure. During the helical intermediate formation (conformers T1 to T3), the intra H-bonds increased. In contrast, the inter H-bonds increased during the *β*-sheet transition (conformer T4→T6).

A similar interaction was observed at the N-terminal of P4 that transitioned from a coil to a 3_10_-helix (T5→T6). Intra-HB started near N-terminal in most of the peptides. In fact, H-bonds between amino acid residues L(5) and R(7) were found in all peptides in different conformers (their occurrence is indicated by small arrows and parenthetic (5,7) notation in (Figure 7). These observations suggested strong residue-residue interactions near the N-termini of peptides. The strong interaction between residues near N-terminal could result from electrostatic attraction between negatively charged aspartic acid (4) and positively charged arginine (7). The strong electrostatic interaction induced H-bonds between L(5) and R(7) leading to initiation of 3_10_-helices at N-termini of peptides. Whereas, intra-HBs increased during the helical intermediate formation (T1→T3), inter-HBs increased during *β*-sheet transitions in the helical intermediate (T4→T6). Since the total number of H-bonds increased from low stability conformer T1 to most stable conformer T6, it suggested the stabilization of conformers by hydrogen bonding. The increase in the total number of H-bonds from the low stability conformer T1 (unaggregated and fluctuating random coils) to the most stable conformer T6 (*β*-sheet dominated aggregate) clearly demonstrated stabilization by hydrogen bonding.

This *structural transition hypothesis*, centred on the evolution of H-bonding networks during aggregation, was further substantiated by a careful inspection of H-bonds in each of the trajectories from both unconstrained and ABF simulations. Most of the peptides showed short transient helices in the early stages of aggregation. Figure 8 clearly shows these transitions in snapshots captured from the early stages of the S1A2P9 aggregation.

**Fig. 8.**
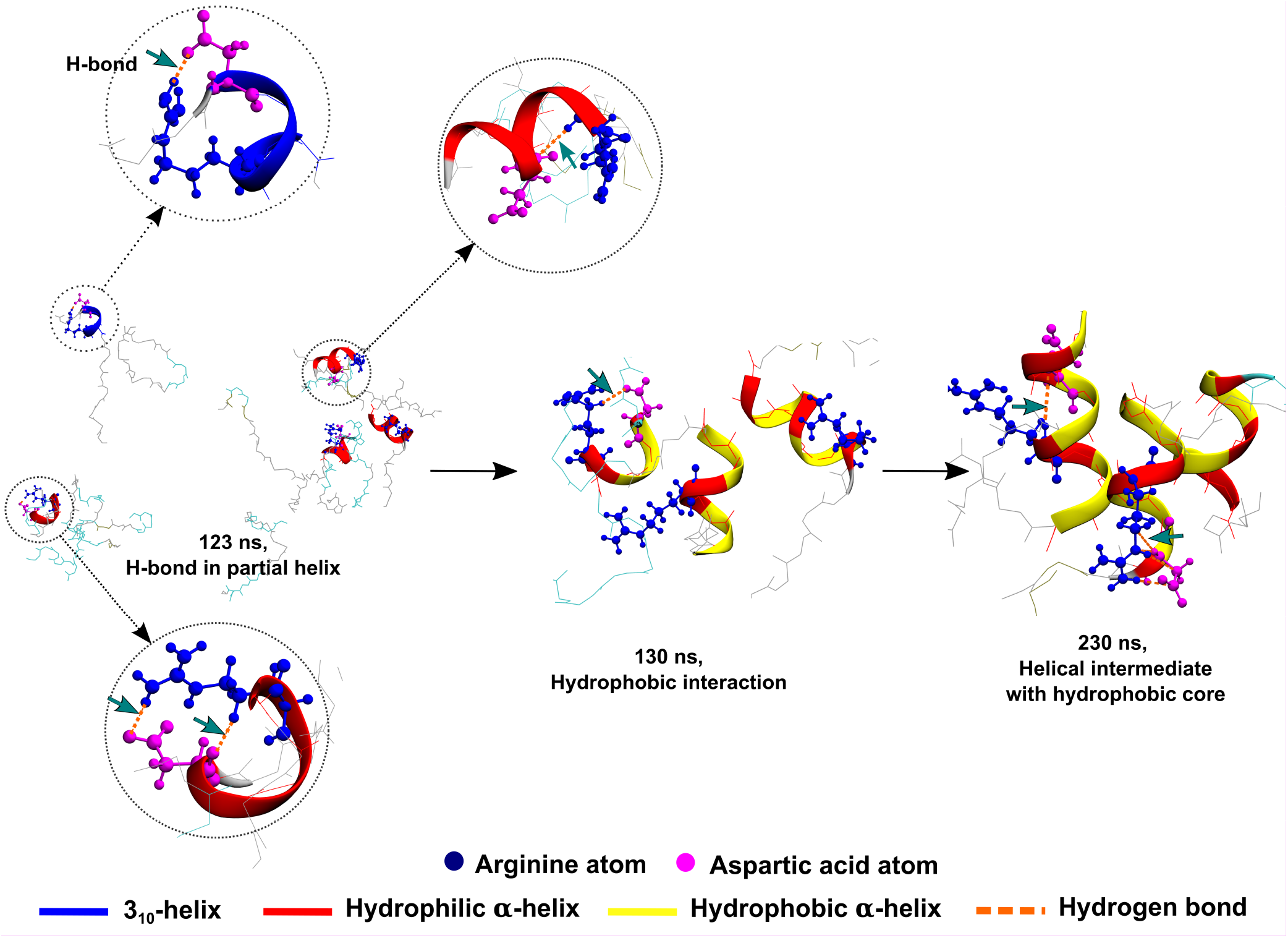
Snapshots of early stages of S1A2P9 aggregate formation. At 120 ns, the electrostatic interaction between negative charged aspartate and positive charged arginine triggered the H-bonds and subsequently partial helix formations in peptides. The hydrophobic interactions started in hydrophobic surfaces (yellow) of peptides at 130 ns. The hydrophobic interaction of partial helical peptides pulled together to form a trimer containing hydrophobic core at 230 ns that further associated with other peptides to form S1A2P9 aggregate.

This result reveals that the electrostatic interaction between aspartate (4) and arginine (7) makes helix prone segment toward N-terminal of the U3.5 peptide. The helical N-terminal of U3.5 peptides plays an important role in hydrophobic interactions to initiate the formation of the helical intermediates. The helical intermediates facilitate the *β*-sheet transitions by increasing local peptide concentration and therefore peptide-peptide interactions.

At 123 ns, electrostatic attraction between D(4) and R(7) residues induced H-bonds in short segments which triggers the formation of short helices. Several unstable short 3_10_-helices were observed before transformation to *α*-helices. The formation of amphipathic *α*-helices leads to greater hydrophobic interactions between different peptides in partial *α*-helical conformations. At 130 ns, the hydrophobic surfaces (yellow) of three peptides (in partial *α*-helical conformations) were primarily driven by hydrophobic interactions that led to the formation of the stable helical trimer. The snapshot at 230 ns showed a stable helical trimer with a hydrophobic core and an enhanced *α*-helical content (compared to 130 ns). Additional random coil peptides attached to the trimeric aggregate via hydrophobic interactions resulted in an increase in the aggregate size. The further evolution of the aggregate structure is also seen in Figure 3 where the additional peptides join the aggregate to form a larger cluster.

### 3.3 Conclusions

Amyloid deposition in tissues is associated with more than fifty human diseases. However, molecular mechanisms of amyloid formation, especially at early stages, are not well understood. Previous studies have revealed formation of transient helical intermediates before mature fibrillation of peptides or proteins into amyloids. In experiments of U3.5 aggregation in water-TFE mixtures, Calabrese *et al*. showed that low amounts (10 – 20% v/v) of TFE (an *α*-helix promoter) increased *β*-sheet aggregation. In contrast, high amounts of TFE (50% v/v) inhibited the *β*-sheet aggregation of U3.5 peptides.

In the current study, microsecond-scale MD simulations were utilized to investigate mechanisms of helical intermediates formation and their role in amyloid formation in antimicrobial, amyloidogenic U3.5 peptides. In conjunction with MD simulations, ABF calculations, provided novel insight into the formation of helical intermediates during aggregation, and their further evolution into *β*-sheet-rich aggregates. From the simulations, a clear picture of peptide aggregation into *β*-sheet dominated aggregates emerged that centred on two important factors; the evolution of *α*-helical intermediates and the critical role of local peptide concentration inside aggregates.

An exhaustive analysis of intermolecular interactions revealed that electrostatic attraction between the negatively charged aspartic acid and positively charged arginine, situated at the 4th and 7th positions respectively from N-terminal, triggered helix formation in U3.5 peptides (Figure 8). Hydrogen bonding between D4 and R7 led to formation of 3_10_-helices at segment 4-DLIR-7 near N-terminus of peptides. Careful analysis of structural transitions showed the 3_10_-helix to be precursors of *α*-helix. Consequently, the emergence of a majority of the 3_10_-helices at the N-terminus, resulted in the N-termini region of peptides being more prone to *α*-helix formation. In contrast, most *β*-sheets formed near the C-termini (Figure 6).

Partial helices that formed in peptides were amphipathic and attraction between hydrophobic surfaces of partial helical peptides led to their aggregation into helical intermediates. As aggregation continued, an increase in local peptide density facilitated the transition of helical intermediates to *β*-sheet dominated aggregates (Figure 3). A correlation analysis between initial *α*-helix content and final secondary structures in the unconstrained simulations revealed that the peptides with low partial helix easily transformed into *β*-sheet. In contrast, peptides with high initial helical content which had more stable helices that were resistant to transformation to *β*-sheets (Figure 2b).

The entire mechanism of the structural transition hypothesis is summarised by Figure 8 and conveys the following sequence of events. Electrostatic attraction between oppositely charged residues at the N-terminus trigger H-bond formation and helix formation. Helical peptides are more stable against thermal fluctuations and aggregate together, driven by hydrophobic interactions, to form helical intermediates. These helical intermediates provide additional stability to the peptide cluster which grows in size (number of peptides). This leads to an increase in local peptide density which enhances peptide-peptide interactions, and finally induces an *α*-helix to *β*-sheet transition in the aggregate.

U3.5 is an amyloid-forming antimicrobial peptide. Therefore, it can be used as a model peptide to study both amyloid formation and antimicrobial drug development. The roles of helical intermediates in *β*-sheet-rich aggregation could be useful in the understanding of the amyloid formation and therapeutic development of amyloidosis. Furthermore, a study of toxic nature of helical intermediate can be useful to develop novel antibiotics against the bacteria.

## Acknowledgement

The authors acknowledge financial supports of Department of Biotechnology (DBT), Government of India (GOI) for project (IMURA0781) at IITB-Monash research academy.

## Notes and references

1 F. Chiti and C. M. Dobson, Annual review of biochemistry, 2017, 86, 27–68.

2 M. D. Benson, J. N. Buxbaum, D. S. Eisenberg, G. Merlini, M. J. Saraiva, Y. Sekijima, J. D. Sipe and P. Westermark, Amyloid, 2018, 25, 215–219.

3 D. J. Selkoe, Neuron, 1991, 6, 487–498.

4 F. Chiti, C. M. Dobson et al., Annual review of biochemistry, 2006, 75, 333–366.

5 A. Gladytz, E. Lugovoy, A. Charvat, T. Häupl, K. Siefermann and B. Abel, Physical Chemistry Chemical Physics, 2015, 17, 918–927.

6 P. T. Lansbury and H. A. Lashuel, Nature, 2006, 443, 774–779.

7 M. D. Kirkitadze, M. M. Condron and D. B. Teplow, Journal of molecular biology, 2001, 312, 1103–1119.

8 J. A. Williamson and A. D. Miranker, Protein Science, 2007, 16, 110–117.

9 T. John, A. Gladytz, C. Kubeil, L. L. Martin, H. J. Risselada and B. Abel, Nanoscale, 2018, 10, 20894–20913.

10 Y. Fezoui and D. B. Teplow, Journal of Biological Chemistry, 2002, 277, 36948–36954.

11 B. Kim, T. D. Do, E. Y. Hayden, D. B. Teplow, M. T. Bowers and J.-E. Shea, The Journal of Physical Chemistry B, 2016, 120, 5874–5883.

12 M. R. Chapman, L. S. Robinson, J. S. Pinkner, R. Roth, J. Heuser, M. Hammar, S. Normark and S. J. Hultgren, Science, 2002, 295, 851–855.

13 C. B. Ramsook, C. Tan, M. C. Garcia, R. Fung, G. Soybelman, R. Henry, A. Litewka, S. O’Meally, H. N. Otoo, R. A. Khalaf et al., Eukaryotic cell, 2010, 9, 393–404.

14 A. M. Bradford, J. H. Bowie, M. J. Tyler and J. C. Wallace, Australian journal of chemistry, 1996, 49, 1325–1331.

15 N. Salinas, E. Tayeb-Fligelman, M. D. Sammito, D. Bloch, R. Jelinek, D. Noy, I. Usón and M. Landau, Proceedings of the National Academy of Sciences, 2021, 118, e2014442118.

16 S. Ray, S. Holden, L. L. Martin and A. S. Panwar, Peptide Science, 2019, 111, e24120.

17 S. Ray, S. Holden, A. K. Prasad, L. L. Martin and A. S. Panwar, The Journal of Physical Chemistry B, 2020, 124, 11659–11670.

18 T. John, T. J. Dealey, N. P. Gray, N. A. Patil, M. A. Hossain, B. Abel, J. A. Carver, Y. Hong and L. L. Martin, Biochemistry, 2019, 58, 3656–3668.

19 A. K. Prasad, C. Tiwari, S. Ray, S. Holden, D. A. Armstrong, K. J. Rosengren, A. Rodger, A. S. Panwar and L. L. Martin, ChemPlusChem, 2022, 87, e202100408.

20 A. Calabrese, Y. Liu, T. Wang, I. Musgrave, T. Pukala, R. Tabor, L. Martin, J. Carver and J. Bowie, 2016.

21 B. Urbanc, M. Betnel, L. Cruz, G. Bitan and D. B. Teplow, Journal of the American Chemical Society, 2010, 132, 4266–4280.

22 J. A. Lemkul and D. R. Bevan, The Journal of Physical Chemistry B, 2010, 114, 1652–1660.

23 T. Takeda and D. K. Klimov, Biophysical journal, 2009, 96, 4428–4437.

24 S. Bhattacharya, L. Xu and D. Thompson, ACS chemical neuroscience, 2019, 10, 2830–2842.

25 Y.-S. Lin, G. R. Bowman, K. A. Beauchamp and V. S. Pande, Biophysical journal, 2012, 102, 315–324.

26 A. V. Rojas, A. Liwo and H. A. Scheraga, The Journal of Physical Chemistry B, 2011, 115, 12978–12983.

27 W. Humphrey, A. Dalke and K. Schulten, Journal of molecular graphics, 1996, 14, 33–38.

28 J. C. Phillips, R. Braun, W. Wang, J. Gumbart, E. Tajkhorshid, E. Villa, C. Chipot, R. D. Skeel, L. Kale and K. Schulten, Journal of computational chemistry, 2005, 26, 1781–1802.

29 U. Essmann, L. Perera, M. L. Berkowitz, T. Darden, H. Lee and L. G. Pedersen, The Journal of chemical physics, 1995, 103, 8577–8593.

30 G. J. Martyna, D. J. Tobias and M. L. Klein, The Journal of chemical physics, 1994, 101, 4177–4189.

31 S. E. Feller, Y. Zhang, R. W. Pastor and B. R. Brooks, The Journal of chemical physics, 1995, 103, 4613–4621.

32 E. Neria, S. Fischer and M. Karplus, The Journal of chemical physics, 1996, 105, 1902–1921.

33 L. Martínez, R. Andrade, E. G. Birgin and J. M. Martínez, Journal of computational chemistry, 2009, 30, 2157–2164.

34 A. Laio and M. Parrinello, Proceedings of the National Academy of Sciences, 2002, 99, 12562– 12566.

35 G. Fiorin, M. L. Klein andJ. Hénin, Molecular Physics, 2013, 111, 3345–3362.

36 H. K. Ho, M. J. Kuiper and R. Kotagiri, Bioinformatics, 2008, 24, 2934–2935.

37 G. E. Crooks, G. Hon, J.-M. Chandonia and S. E. Brenner, Genome research, 2004, 14, 1188– 1190.

38 R. Tycko, Protein Science, 2014, 23, 1528–1539.

39 A. Abedini and D. P. Raleigh, Protein Engineering, Design & Selection, 2009, 22, 453–459.

40 M. Fändrich, Journal of molecular biology, 2012, 421, 427–440.

41 A. L. Serrano, J. P. Lomont, L.-H. Tu, D. P. Raleigh and M. T. Zanni, Journal of the American Chemical Society, 2017, 139, 16748–16758.

42 R. Armen, D. O. Alonso and V. Daggett, Protein Science, 2003, 12, 1145–1157.

43 C. A. Rohl and A. J. Doig, Protein science, 1996, 5, 1687–1696.

44 Z. Bikadi, L. Demko and E. Hazai, Archives of Biochemistry and Biophysics, 2007, 461, 225– 234.

