## Supplementary information for "Helical Intermediate Formation and Its Role in Amyloid of an Amphibian Antimicrobial Peptide^†^"

### S-I ABBREVIATIONS

U3.5, uperin 3.5;

MD, molecular dynamic;

AMP, antimicrobial peptide;

CD, circular dichroism;

IR, infrared spectroscopy;

RMSD, root mean square distance;

$R_g$, radius of gyration

SASA, solvent accessible of surface area

SDS, sodium dodecyl sulfate;

TFE, 2,2,2-trifluoro- ethanol;

DOPE, (1,2-dioleoyl-sn-glycero-3-phosphoethanolamine)

DOPG, (1,2-dioleoyl-sn-glycero-3-phospho-(1′-rac-glycerol))

ThT, thioflavin T;

ABF, adaptive biasing force;

PMF, potential of mean force;

HB, hydrogen bond;

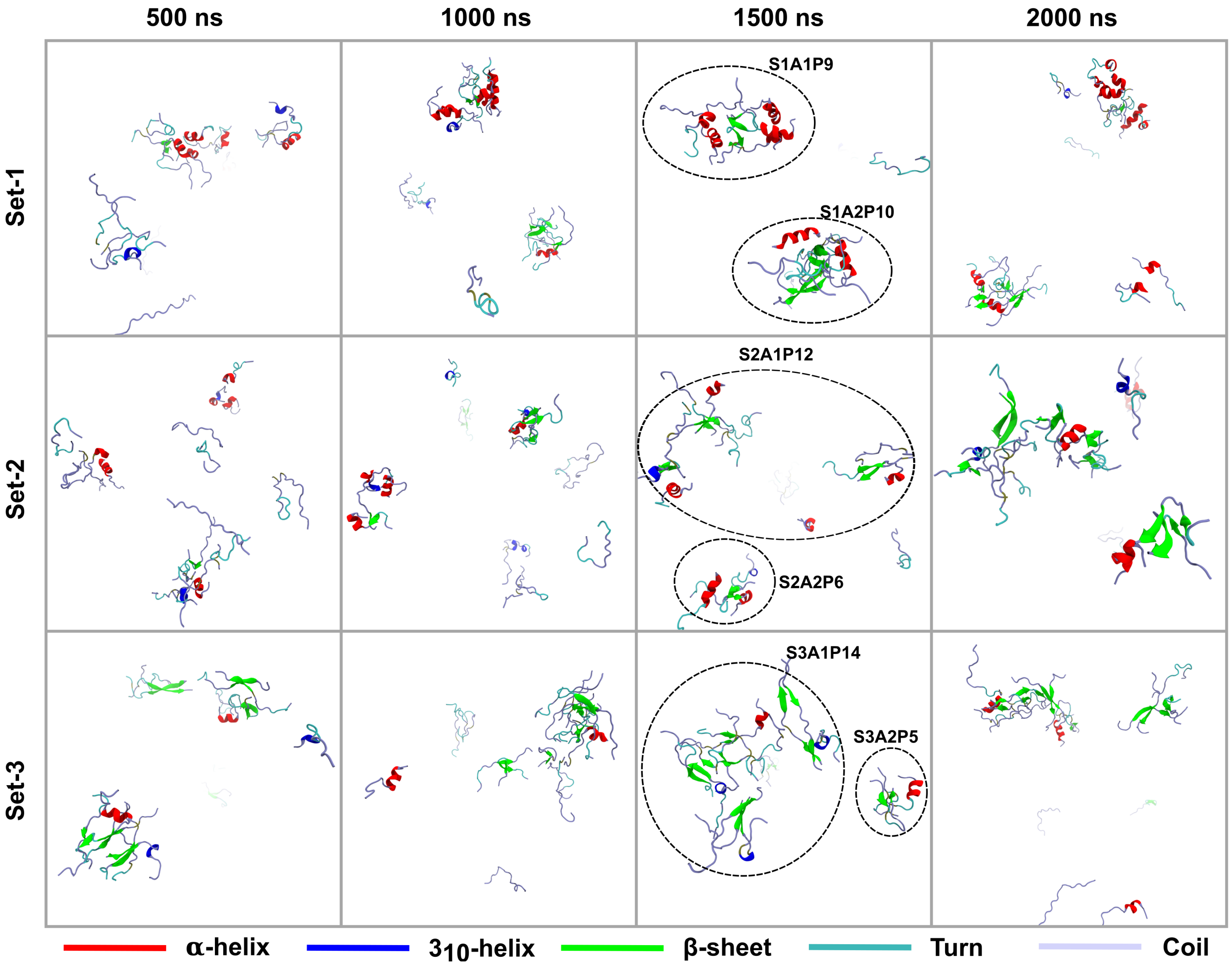


**Fig. S1** Snapshots of the Unconstrained simulation, captured at different times for each trajectory, are shown in the figure. The simulations were started with twenty unaggregated U3.5 peptides, containing random coil or partial helical structures, that turn into $\beta$-sheet-rich aggregates. The aggregates are shown by round circle at 1500 ns.

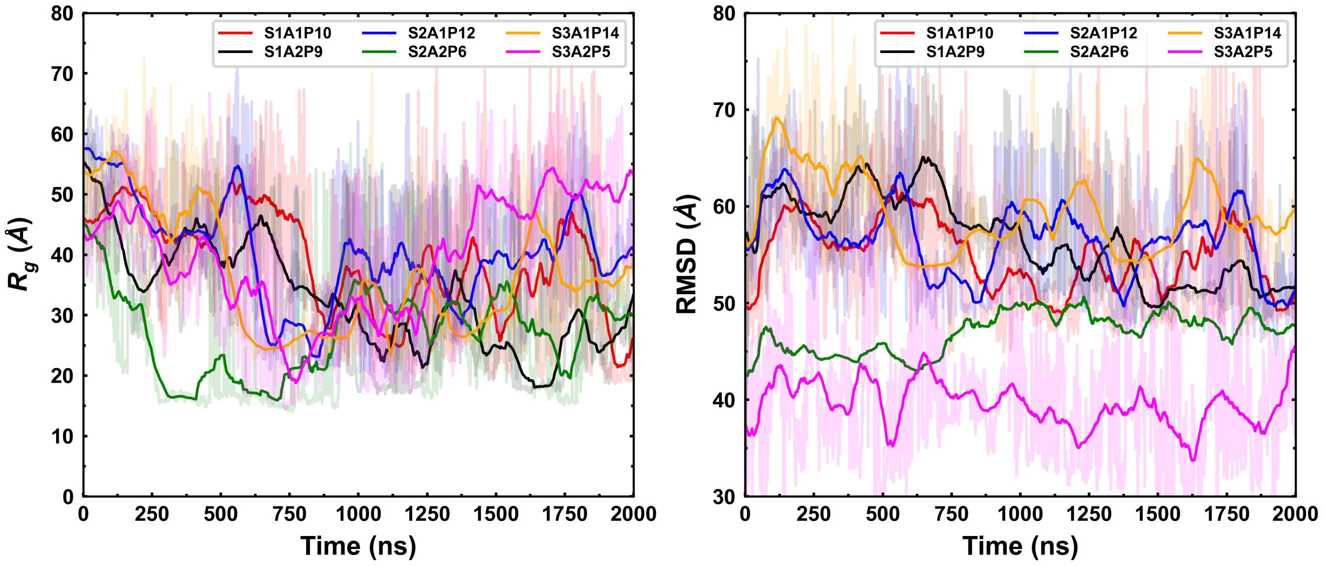


**Fig. S2** Radius of gyration ($R_g$) and root mean square deviation (RMSD) of each aggregate observed in unconstrained simulations are shown in the figure (running averaged with 100 data points). The radius of gyration of aggregates was decreased significantly till $\approx$ 900 ns, since peptides were assembled in aggregates during this period, which remained stable over the end of the simulations (except S3A2P5 aggregate). The initial structures were used to calculate the respective RMSD of aggregates. The fluctuation of RMSD indicated structural changes in corresponding aggregates.

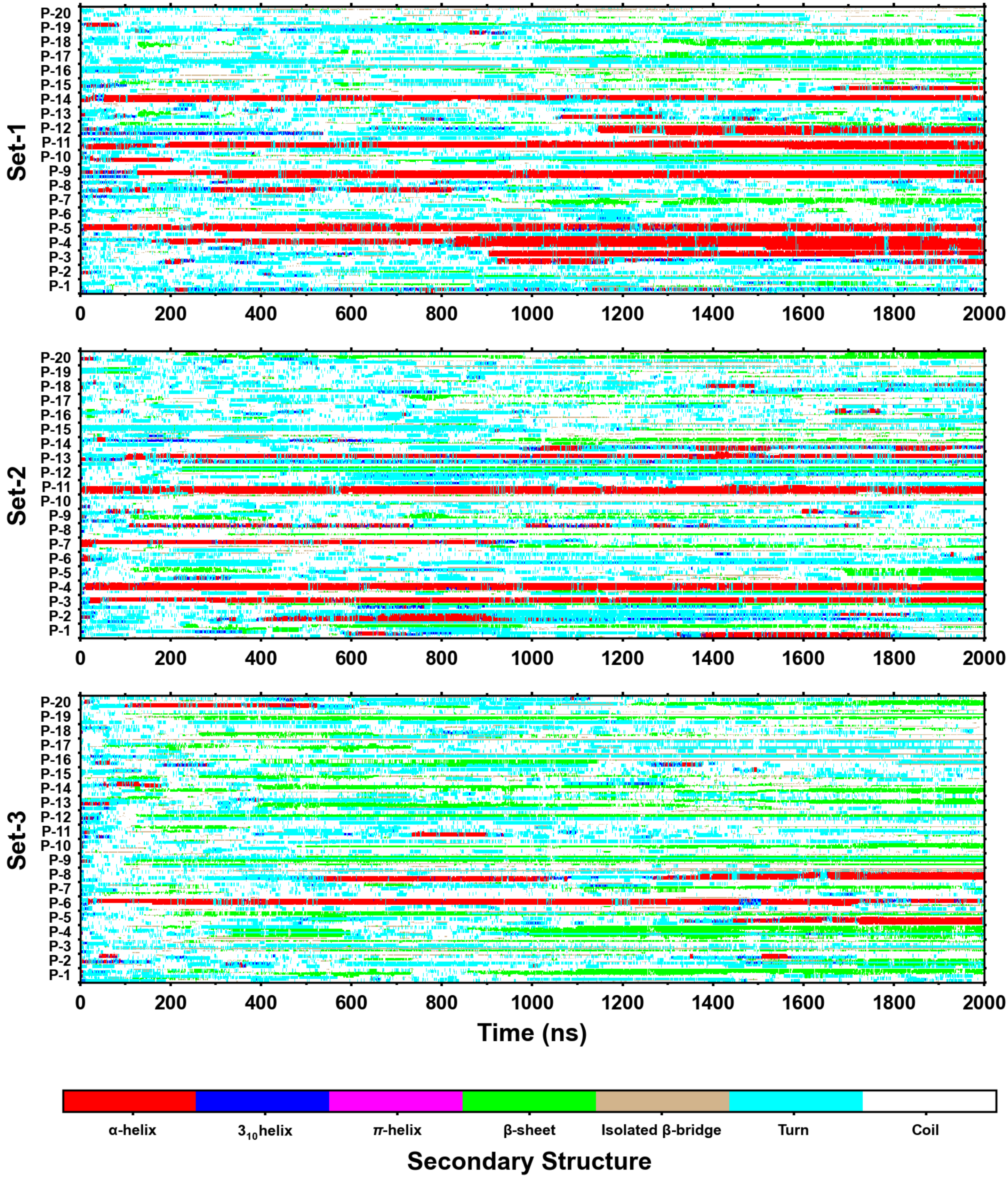


**Fig. S3** Heat map of secondary structures for all trajectories of unconstrained simulations. The y-axis of the heat map is peptides, while the x-axis is simulation time. The length of color strip can indicate the stability of the corresponding secondary structure. $3_{10}$-helices appeared, mostly, where the $\alpha$-helices are formed or vanished.

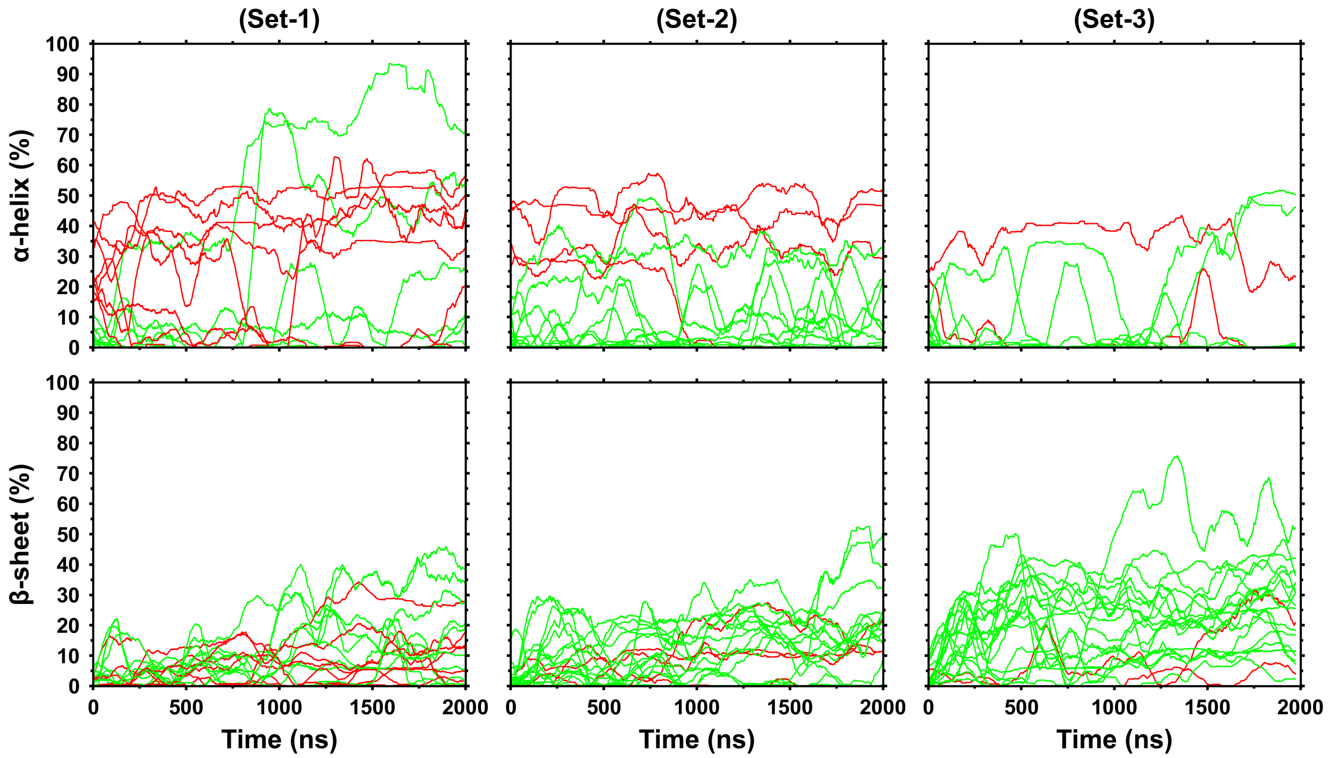


**Fig. S4** Secondary structure evolution for individual peptides of the unconstrained simulations. Each line represents an individual peptide, while the color is based on its initial helical content. The red line corresponds to peptides if initial helical content $> 15\%$, and all else belongs to the green line.

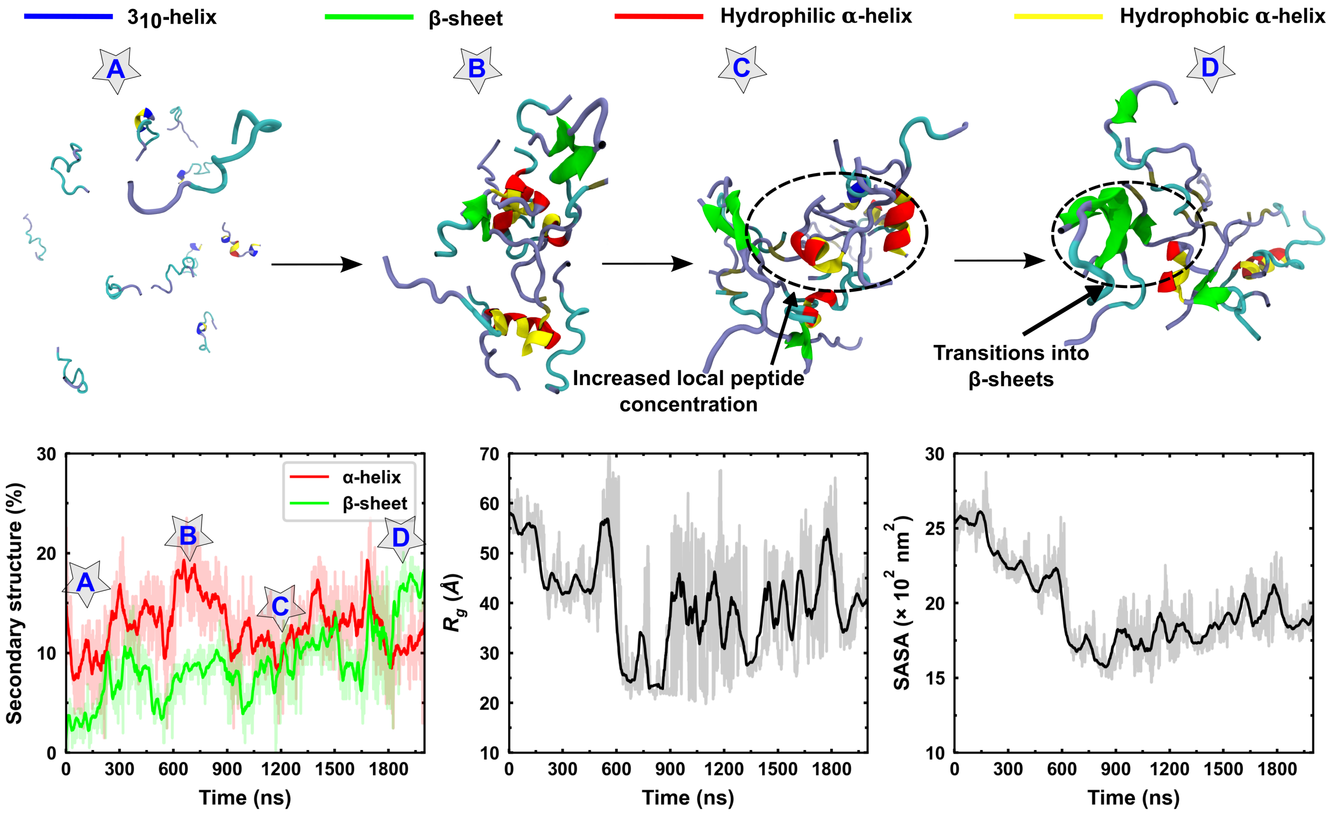


**Fig. S5** The snapshots correspond to different stages of the S2A1P12 aggregate formation are shown along with the secondary structure, $R_g$, and SASA. The aggregate formation started with the clustering of unaggregated peptides, containing mostly random coil structures (conformer A), that led to the formation of helical intermediate (conformer B). During A⟶B transitions, the $R_g$ decreased with a significant amount of helix formations. Peptides in aggregate were rearranged which led to the formation of increased local peptide concentration (conformer C). Once the peptide-peptide interaction was enhanced by the increased local peptide concentration, the $\beta$-sheets formed in aggregate (conformer D).

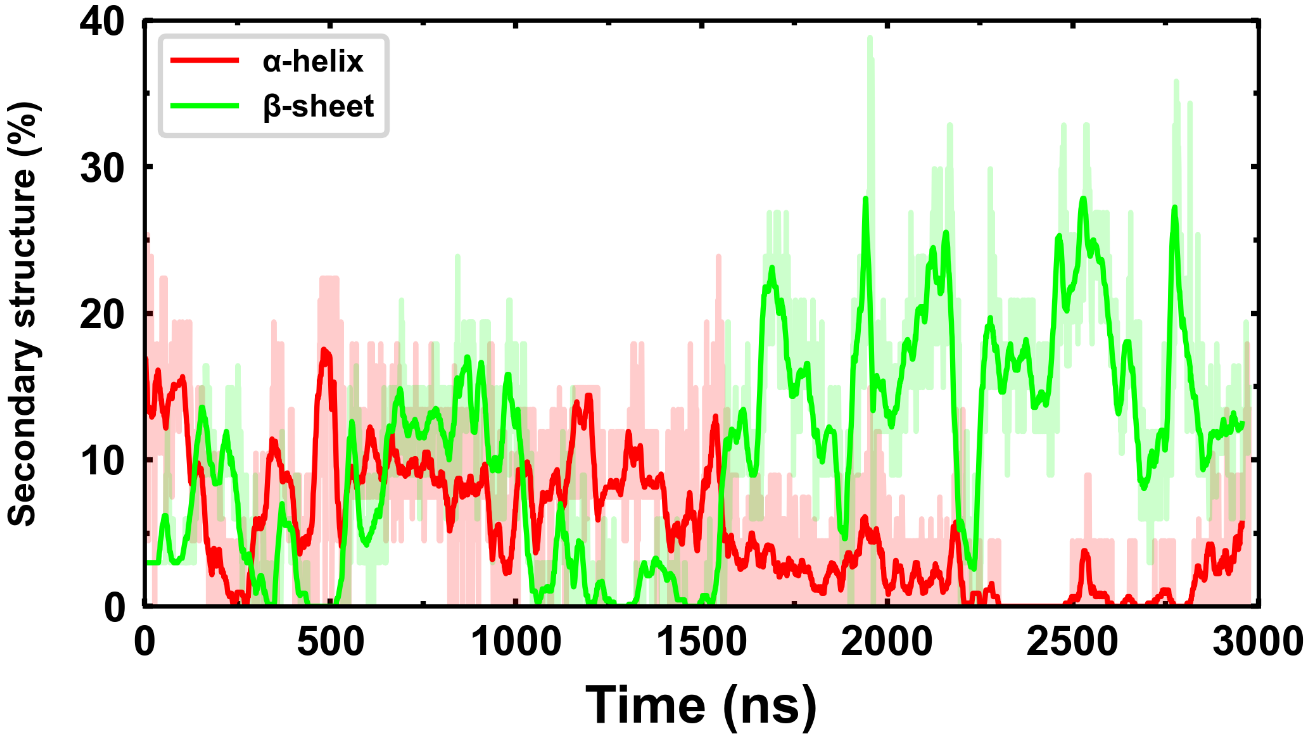


**Fig. S6** Secondary structure evolution of ABF simulation. The $\beta$-sheet content in peptides was continuously increased with the loss of helical content of peptides

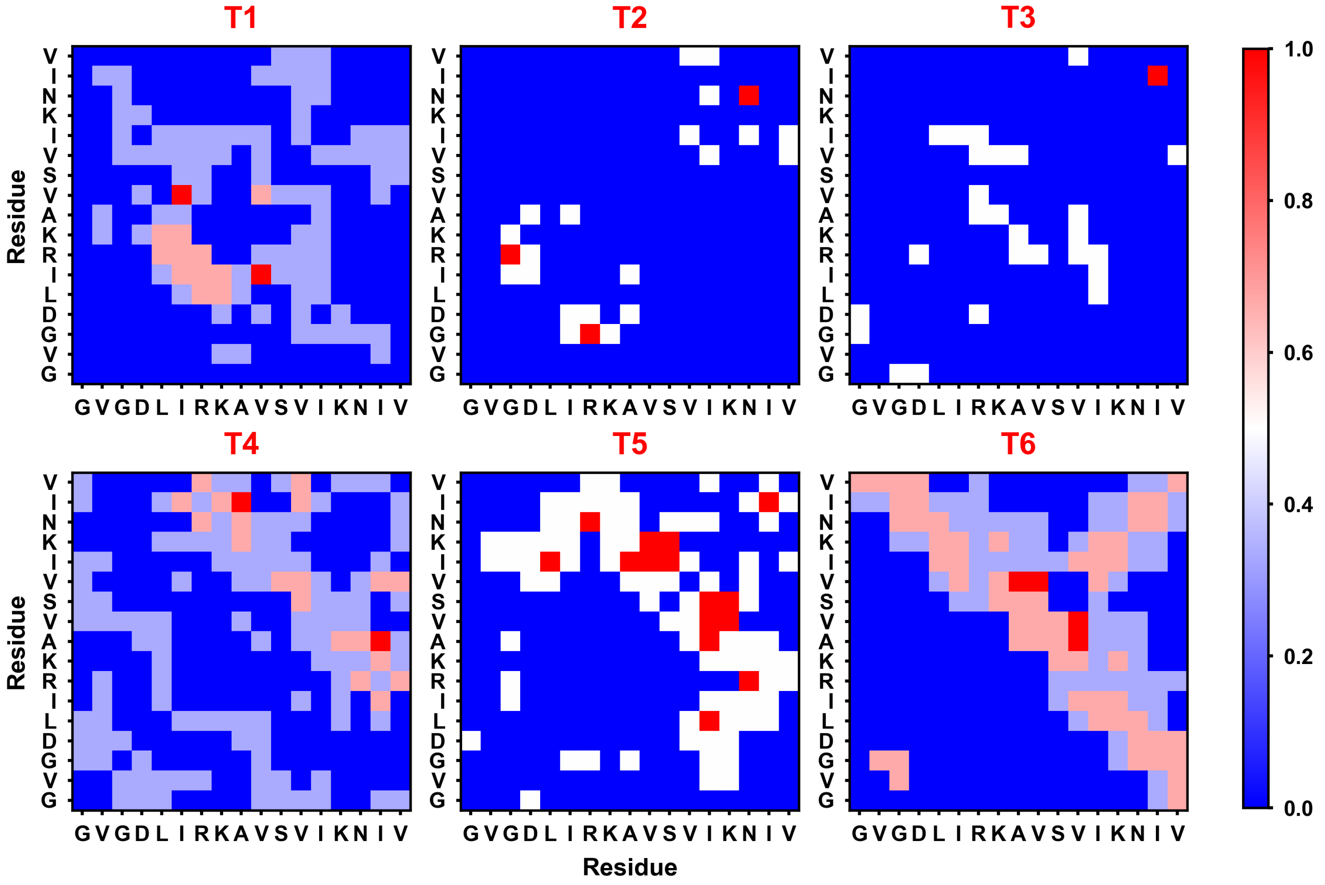


**Fig. S7** Inter-peptide contact map for individual conformer of ABF simulation are shown, the blue to red color spectrum shows lowest to highest contact. During helical intermediate formation (T1⟶T3), the N-terminus of peptides showed strong interactions (specifically, aspartate (4) and arginine (7) residues). Once the helical intermediate formed, the peptide-peptide interactions toward the C-terminus became dominant (cross-diagonal contacts indicate the antiparallel $\beta$-sheet structure)

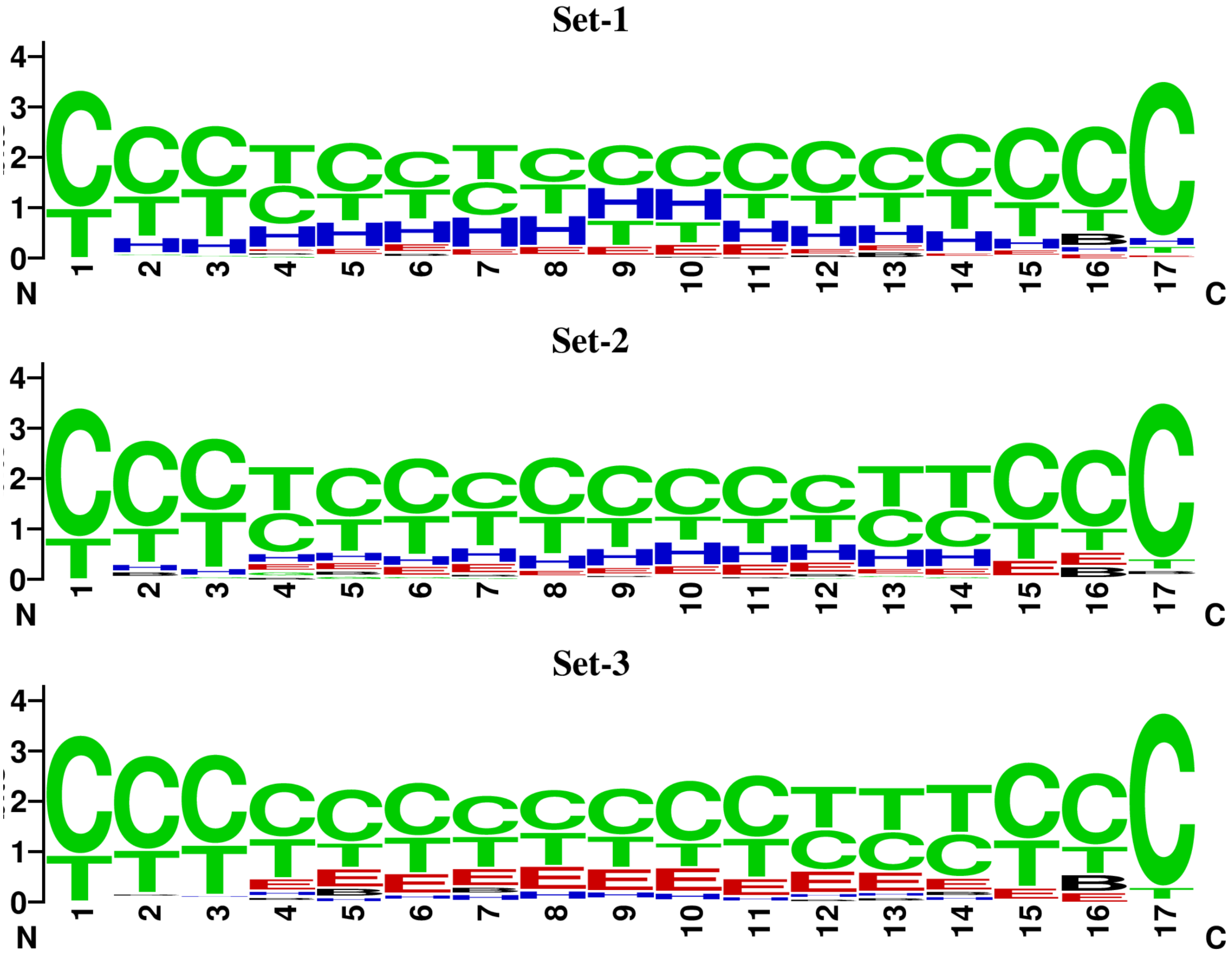


**Fig. S8** Sequence logo of the secondary structure represents the relative frequency of structural components of each residue. Symbols (C = coil, T = turn, H = helix, E = $\beta$-sheets, and B = $\beta$-bridge) are used for the secondary structure components. The height of the symbol indicates the relative frequency of the secondary structure for the corresponding residue. The sequence logos were calculated using the structural information of 10 ns window with 100 ns gap over trajectories of unconstrained simulations.
